# Chromosomal instability promotes cell migration and invasion via EFEMP1 secretion into extracellular vesicles

**DOI:** 10.1101/2024.10.21.619397

**Authors:** Siqi Zheng, Ruifang Tian, Yuanyuan Liu, Rene Wardenaar, Marjan Shirzai, Laura Kempe, Emma Dijkstra, Eliza Warszawik, Maria Suarez Peredo Rodriguez, Klaas Sjollema, Petra L. Bakker, Patrick van Rijn, Michaela Borghesan, Judith Paridaen, Stefano Santaguida, Floris Foijer

## Abstract

Triple-negative breast cancer (TNBC) is characterised by high rates of chromosomal instability (CIN) and a tumour microenvironment (TME) modulated by extracellular vesicles (EVs). To assess how CIN might affect the TME in TNBC, we studied the EV landscape of TNBC cell lines with induced CIN. We find that CIN leads to increased secretion of EVs and that these EVs promote cell migration of recipient cells. EVs are enriched for extracellular matrix (ECM) proteins, including EFEMP1. Indeed, modulation of EFEMP1 levels in EVs significantly alters migration behaviour of EV-treated cells. We show that EFEMP1 expression is regulated by STAT1, and that EVs from STAT1-deficient cells no longer promote migration, which can be rescued by overexpression of EFEMP1 in STAT1-null cells. Xenografting TNBC cells with EFEMP1 enriched cells promotes migration in zebrafish embryos, suggesting that EFEMP1 expression is a factor that promotes metastasis. Together our results uncover a novel role for CIN in shaping the TME of TNBC and identify EFEMP1 as a potential therapeutic target to prevent cell migration within the TME.

## Introduction

Triple-negative breast cancer (TNBC) is defined by the loss of the estrogen receptor (ER), progesterone receptor, and human epidermal growth factor 2 (HER2) and is associated with poor prognosis (Dietze *et al*, 2015). TNBC frequently displays chromosomal instability (CIN), an increased frequency of chromosome segregation errors during mitosis (Bianchini *et al*, 2016; Vargas-Rondón *et al*, 2017). CIN in TNBC is associated with cancer cell evolution, immune evasion, an altered tumour microenvironment (TME), increased metastasis and thus, a poor outcome (Hoevenaar et al, 2020; Gao et al, 2016; Bakhoum & Cantley, 2018; Li et al, 2021b).

Extracellular vesicles (EVs), such as exosomes and microvesicles secreted by cancer cells and other cells in the TME, are known to play an important role in the communication between cancer cells and the TME, thus shaping the TME (Bhome *et al*, 2022). Indeed, EVs, can serve as biomarkers for cancer and genomic instability, e.g. when measured in peripheral blood (Minciacchi *et al*, 2015; Willms *et al*, 2018; Martins *et al*, 2023; Bao *et al*, 2021). Although the contents of various EVs have been extensively studied to identify the roles of different lncRNAs, microRNAs, and proteins in tumour progression (Minciacchi *et al*, 2015; Willms *et al*, 2018; Martins *et al*, 2023; Bao *et al*, 2021), the role of EVs in a chromosomal instable background remains unexplored.

In this study, we investigate how a CIN phenotype in cells influences the secretion and content of EVs and the impact of these EVs on their neighbouring cells. For this, we compare the composition of EVs secreted by CIN^HIGH^ and CIN^LOW^ TNBC cells and the effect of their secreted EVs on CIN^LOW^ TNBC cells. We find that CIN leads to increased secretion of EVs and that EVs secreted from CIN^HIGH^ TNBC cells promote migration and invasion of isogenic CIN^LOW^ TNBC cells. We identify the extracellular matrix protein EFEMP1, also known as Fibulin-3 and a known prognostic factor in TNBC (McHenry & Prosperi, 2023; Noonan *et al*, 2018; Hu *et al*, 2011) to be enriched in EVs secreted by CIN^HIGH^ TNBC cells. The induction of EFEMP1 expression is linked to induced chromosomal instability (CIN) and relies on the presence of Signal Transducer and Activator of Transcription 1 (STAT1), a crucial factor in breast cancer (Banik *et al*, 2021). Overexpression of EFEMP1 rescues the delayed migration observed in TNBC cells lacking STAT1 and restores the invasive potential of EVs originating from STAT1-expressing cells, thereby promoting the invasive phenotype. In conclusion, we have uncovered a CIN-driven pathway involving EFEMP1 and STAT1 that promotes cell migration within the TME through a paracrine mechanism, potentially contributing to the increased metastatic potential of CIN^HIGH^ TNBC.

## Results

### Chromosomal instability promotes extracellular vesicle production and release in TNBC cancer cells

To compare the effects between EVs released by cells with low and high rates of CIN, we use two frequently used and well-characterized TNBC cell lines, BT549 and MDA-MB-231. To induce CIN^HIGH^ phenotypes, we use the MPS1 inhibitor reversine (Santaguida *et al*, 2010), a widely used compound for this purpose (Bosco *et al*, 2018; Hiruma *et al*, 2016; Garribba *et al*, 2023; Ippolito *et al*, 2021). CIN phenotypes were quantified by time-lapse imaging according to established protocols (Crozier *et al*, 2022; Thu *et al*, 2018; Huis In ’t Veld *et al*, 2019), and confirmed that MPS1 inhibition increased the rate of mitotic abnormalities in a dose-dependent manner in both BT549 and MDA-MB-321 TNBC cells (**Fig. 1A**). To determine whether CIN affects EV production and secretion, we first quantified CD63, an established marker of EVs, by immunofluorescence in BT459 and MDA-MB-231 cells (**Fig. 1B, C**) and found that CD63 levels were significantly increased in both BT549 and MDA-MB-231 cells upon the induction of a CIN phenotype (**Fig. 1D, E**). We then purified EVs secreted into the medium by CIN^HIGH^ and CIN^LOW^ BT549 cells by ultracentrifugation and assessed EV shape by transmission electron microscopy, which confirmed their integrity (Théry *et al*, 2018) (**Sup. Fig. 1A**). Western blot analysis for CD63 and CD81 (another EV marker) confirmed that the isolates indeed were EVs (**Fig. 1F**). Calnexin (endoplasmic reticulum) and beta-actin were used as negative controls (Théry *et al*, 2018). To further characterize the EVs, we used nanoparticle tracking (Théry *et al*, 2018), which revealed that induction of CIN significantly increased EV release, both for BT549 cells as well as MDA-MB-231 cells, while EV size remained unaltered with EV sizes ranging from 40-200 nm (**Fig. 1G, Sup. Figure 1B, C**). We conclude that CIN induced by MPS1 inhibition increases the secretion of EVs for BT549 and MDA-MB-231 TNBC cells.

**Figure 1.**
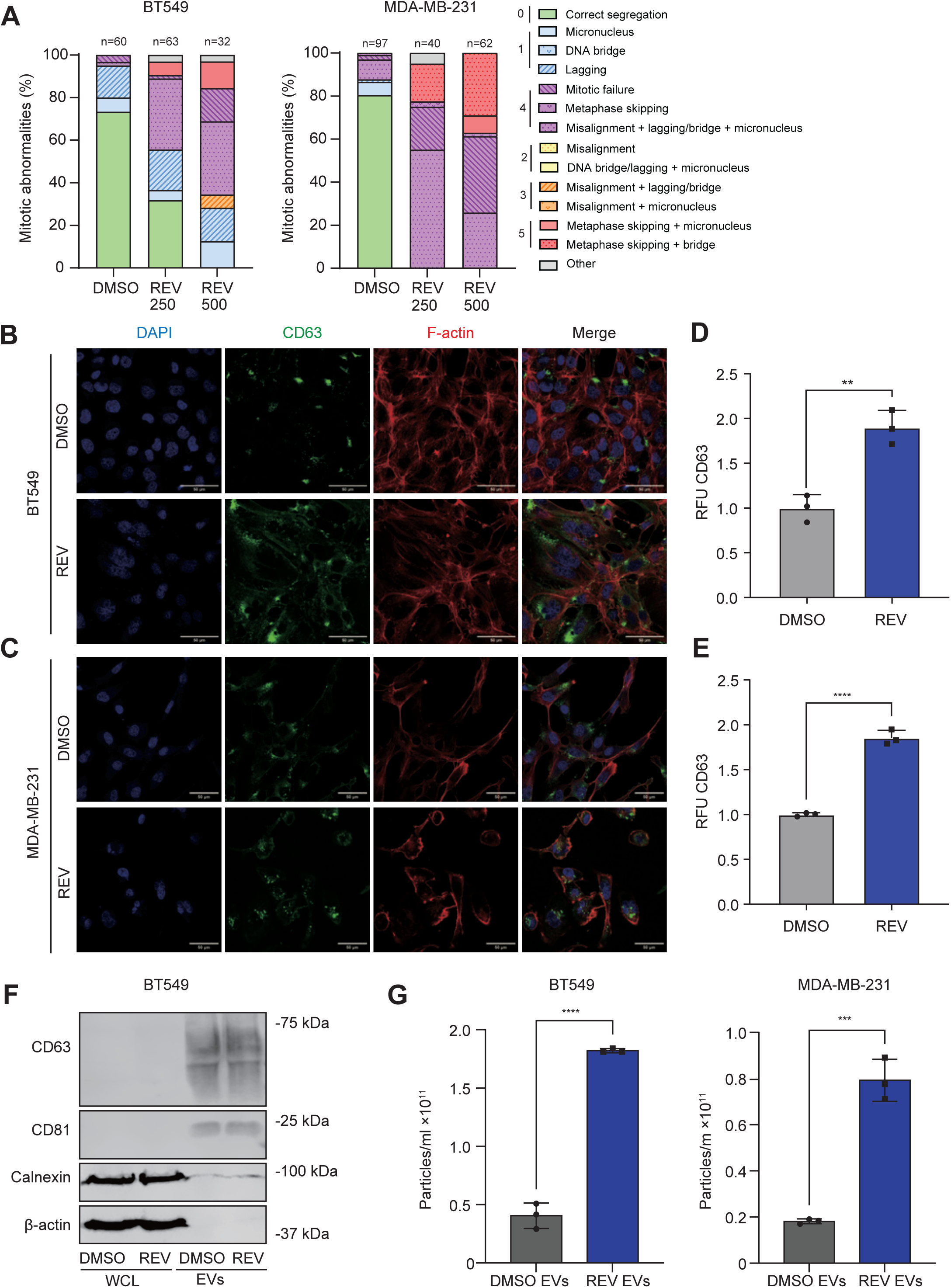
Chromosomal instability promotes extracellular vesicle production and release in TNBC cells. (**A**) Quantitative analysis of mitotic aberrations in BT549 and MDA-MB-231 cells after exposure to 250 nm and 500 nm reversine (REV). (**B, C**) CD63 and F-Actin immunofluorescence labelling in BT549 (**B**) and MDA-MB-231 (**C**) cells treated with 500 nm reversine for 72 hours, compared to DMSO controls. DAPI was used to label nuclei. (**D, E**) Quantification of CD63 immunofluorescence intensity in BT549 (**D**) and MDA-MB-231 (**E**) cells in presence and absence of CIN phenotypes. *p <0.05; **, p <0.01; ***, p-<0.001; ****, p < 0.0001. RFU is relative fluorescence units, REV is reversine. (**F**) Western blot for EV markers for CD63 and CD81.Calnexin and beta-Actin serve as a loading control for whole lysates and negative control for EVs. (**G**) Concentrations of EVs isolated from BT548 (left panel) and MDA-MB-231 (right panel) cells as determined by nanoparticle tracking analysis.

**Supplementary Figure 1.**
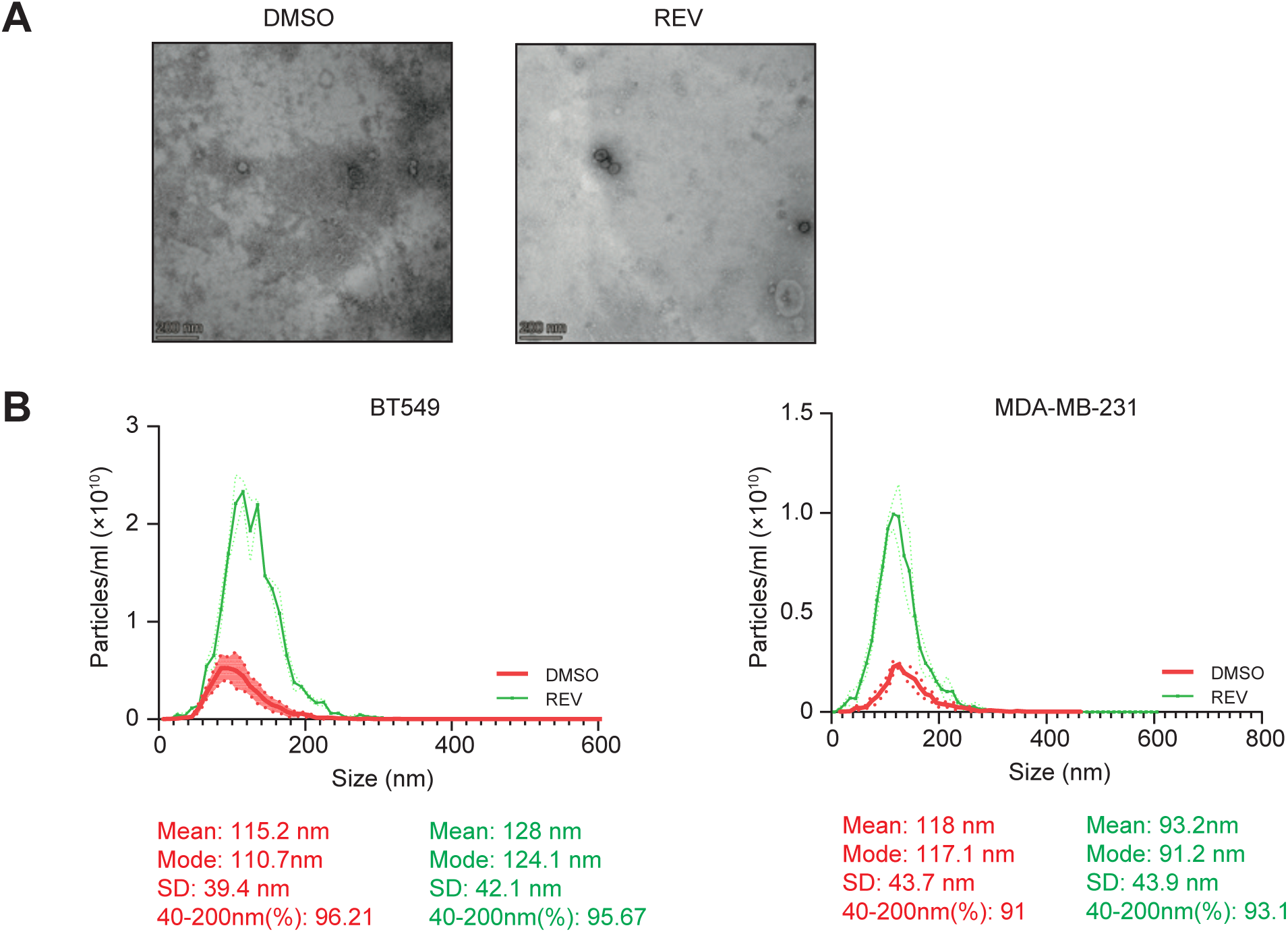
Biophysical characterization of EVs. (**A**) Transmission electron microscopy (TEM) images showing EV morphology when isolated from BT549 treated cells. Scale bar: 200 nm. (**B, C**) Nanoparticle tracking analysis of EVs isolated from BT549 cells (**B**) and MDA-MB-231 cells (**C**), to determine EV size distribution and EV concentrations.

### CIN^HIGH^ TNBC cell-derived EVs promote invasion and migration in a paracrine manner

Next, we investigated the functional impact of EVs released by both CIN^LOW^ and CIN^HIGH^ TNBC cells. We first assessed whether purified EVs are taken up by recipient cells. To achieve this, we incorporated a pHluorin_M153R reporter (Sung *et al*, 2020) into BT549 cells to fluorescently label EVs. We induced CIN phenotypes in these reporter cells, collected labelled EVs, transferred them onto BT549 recipient cells, and monitored the cells via time-lapse imaging. This analysis confirmed the uptake of transferred EVs by the recipient cells (**Sup. Movie 1**). To determine functional consequences of these EVs, we then exposed BT549 cells for 24 hours to EVs isolated from either DMSO-(CIN^LOW^ EVs) or reversine-treated (CIN^HIGH^ EVs) and quantified EdU incorporation as a readout of cell proliferation. We observed no differences in EdU incorporation between cells treated with CIN^LOW^ or CIN^HIGH^ EVs, nor cells that received no EVs, indicating that BT549-isolated EVs and MDA-MB-231-isolated EVs do not influence proliferation of recipient cells. (**Fig 2A, B, Sup. Fig. 2A**). As tumour-derived EVs have previously been associated with increased invasiveness and metastasis (Adams *et al*, 2021; Sun *et al*, 2021), we then determined the impact of BT459 and MDA-MB-321 CIN^LOW^ and CIN^HIGH^ derived EVs on migration and invasion using a trans-well assay in combination with uncoated or Matrigel-coated surfaces (Vasudevan *et al*, 2020). We found a significant increase in migration and invasion when recipient BT549 and MDA-MB-231 cells were treated with CIN^HIGH^ EVs compared to treatment with CIN^LOW^ EVs (**Fig. 2C, D**). In contrast, the cells in which CIN was induced and that were used and from which EVs were isolated showed impaired migration and invasion. This implies that EVs derived from CIN^HIGH^ cells contain factors that facilitate cell migration and invasion in a paracrine manner (**Sup. Fig. 2C, D**).

**Figure 2.**
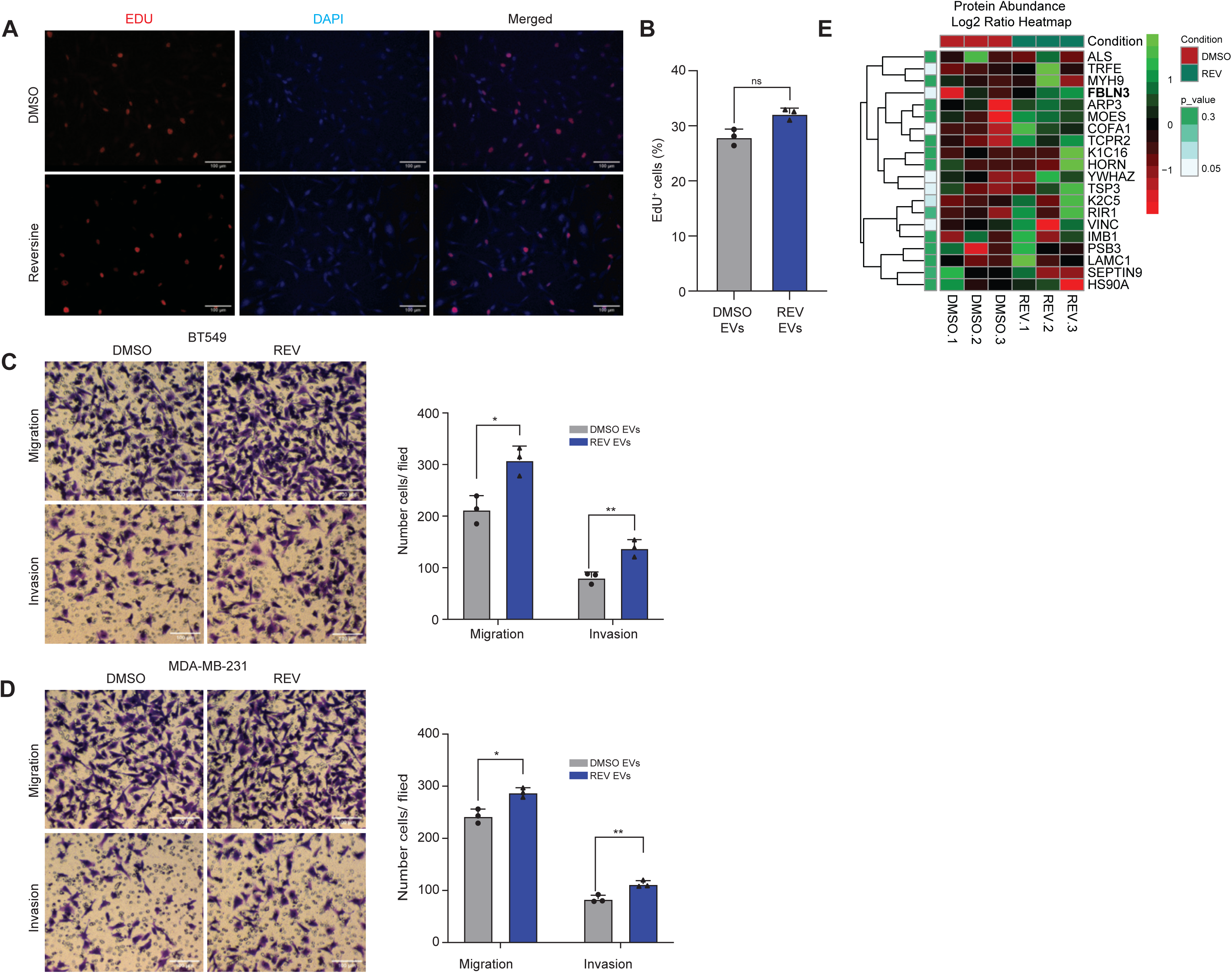
Paracrine effects of EVs isolated from CIN+ BT549 and MDA-MB-231 cells on cell proliferation, migration, and invasion. (**A**) Representative images of EdU incorporation in BT549 cells treated with EVs from DMSO or REV treated cells. Scale bar: 100 µm. (**B**) Quantification of EdU incorporation as shown in (A) for three biological replicates. Statistical analysis was performed using two-sided T-tests (*p < 0.05; **p < 0.01; ***p < 0.001; ****p < 0.0001). (**C, D**) Migration and invasion of BT549 (**C**) and MDA-MB-231 (**D**) cells treated with EVs isolated from BT549 (**C**) or MDA-MB-231 cells (**D**) treated with DMSO (CIN^LOW^) or REV (CIN^HIGH^) using trans-well assays. Statistical significance was determined using two-sided T-tests (*p < 0.05, **p < 0.01, ***p < 0.001, ****p < 0.0001). Experiments were performed as biological triplicates. Scale bar: 100 µm. (**E**) Heatmap of peptides enriched in EVs isolated from BT549 cells treated with REV compared to controls.

To identify factors that mediate the effect on migration, we quantified protein contents of CIN^HIGH^ and CIN^LOW^ BT549 EVs using label-free mass spectrometry (**Sup. Table 1**, **Fig 2E**, most significant peptides). A general gene ontology (GO) analysis showed that the content of our EVs was enriched for ‘exosomes’ (**Sup. Fig. 2B**), validating our EV isolation protocol. Furthermore, ‘Integrin binding’ and ‘Cell adhesion, were among the topmost significantly enriched categories for CIN^HIGH^ EVs in GO and KEGG (Kyoto Encyclopedia of Genes and Genomes) analyses (Luo & Brouwer, 2013; Kanehisa *et al*, 2021; Ge *et al*, 2020) (**Sup. Fig. 2E**). Given its known association with migration and invasion phenotypes (Noonan *et al*, 2018), we decided to further investigate Fibulin3 (FBLN3, **Fig. 2E)**. Fibulin3 is encoded by the *EFEMP1* gene, which is the name we will use in the rest of this study. Indeed, Western blot analysis confirmed that EFEMP1 was present in whole cell lysates as well as EVs, unlike Calnexin and beta-Actin (**Sup Fig. 2F**). We conclude that EVs isolated from CIN^HIGH^ TNBC cells promote migration and invasion in a paracrine manner and the EFEMP1, an EV-enriched factor, is a candidate driver of this phenotype.

**Supplementary Figure 2.**
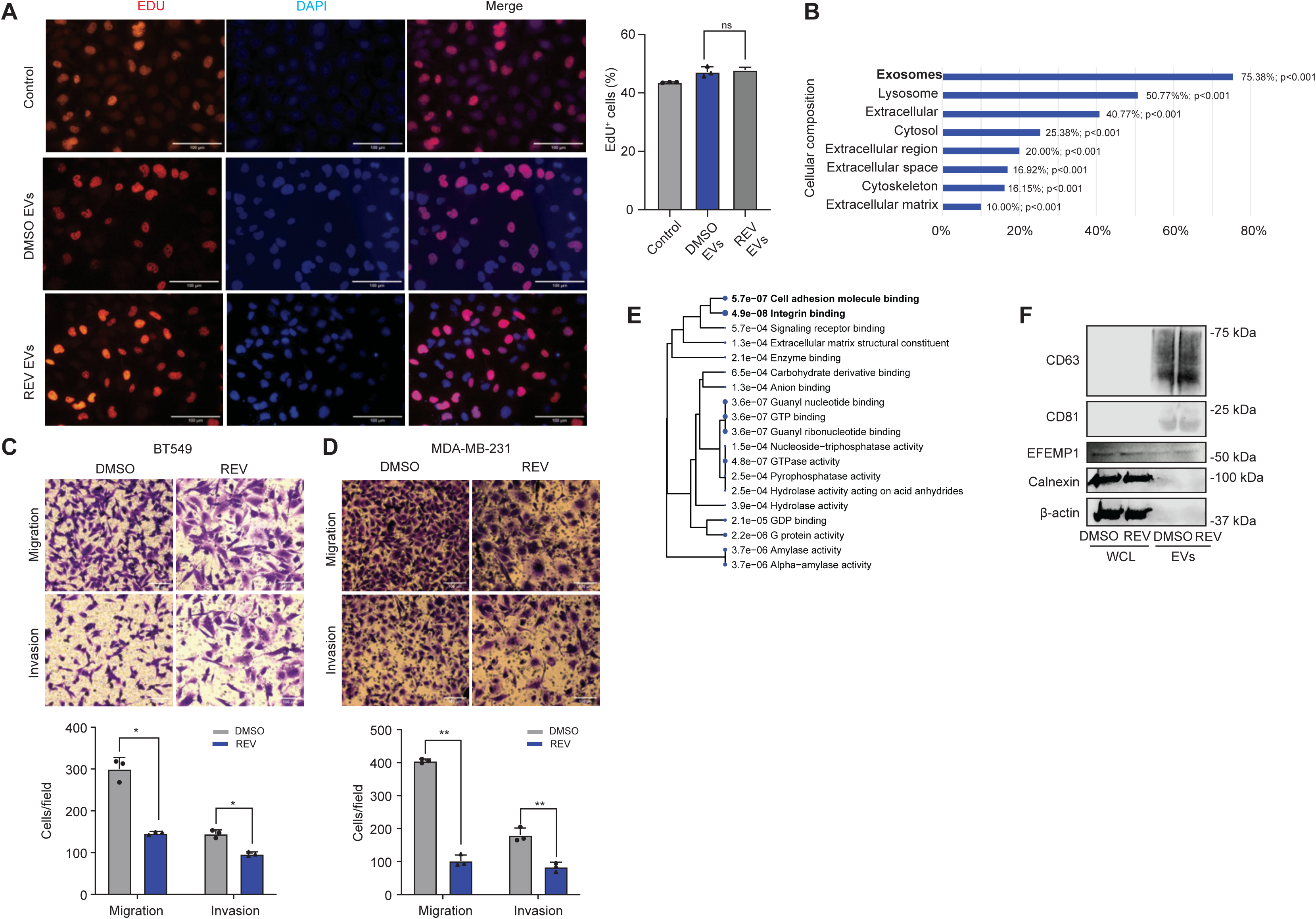
Functional analysis of cellular responses to CIN-derived EVs and Mps1 inhibition in breast cancer cells. (**A**) Representative images of EdU incorporation in MDA-MB-231 cells treated with EVs from DMSO or REV treated cells and quantification for three biological replicates. Statistical analysis was performed using two-sided T-tests (*p < 0.05; **p < 0.01; ***p < 0.001; ****p < 0.0001).) (**B**) Cellular component analysis of differentially expressed proteins using Funrich 3.1.4. software. (**C, D**) Migration and invasion of BT549 (**C**) and MDA-MB-231 (**D**) cells treated with DMSO (CIN^LOW^) or REV (CIN^HIGH^) using trans-well assays. Statistical significance was determined using two-sided T-tests (*p < 0.05, **p < 0.01, ***p < 0.001, ****p < 0.0001). Experiments were performed as biological triplicates. Scale bar: 100 µm. (**E**) Gene Ontology (GO) analysis on the proteins enriched in CIN+ EVs compared to CIN-EVs. (**F**) EFEMP1 levels in BT549 cell lysates and BT549 EVs treated with DMSO (CIN^LOW^) or REV (CIN^HIGH^) detected by Western blot.

### EFEMP1-Enriched EVs promote migration and invasion in recipient TNBC cells

To investigate the role of increased EFEMP1 levels in CIN^HIGH^ EVs in migration and invasion observed in TNBC cells upon EV treatment, we manipulated EFEMP1 expression in BT549 (**Sup. Fig. 3A, B**) and MDA-MB-231 (**Sup. Fig. 3C, D**) in TNBC cells using overexpression and shRNA constructs. We then determined the impact of EFEMP1 modulation on the migratory and invasive potential of these cells directly. Trans-well assays revealed that overexpression of EFEMP1 promotes migration and invasion of both BT549 as well as MDA-MB-231 cells (**Sup. Fig. 3E-H**). Similarly, scratch assays (Jonkman *et al*, 2014) showed that EFEMP1 overexpression promotes invasiveness of BT549 cells while knocking EFEMP1 down reduced invasiveness (**Sup. Fig. 3I, J**). Conversely, downregulation of EFEMP1 impaired migration and invasion of both cell types (**Sup. Fig. 3K-N**). After having evaluated the effects of EFEMP1 expression levels on TNBC cells directly, we assessed the paracrine effect of these cells by isolating EVs of EFEMP1 overexpressing cells (EFEMP1^OEX^ EVs) and knockdown (EFEMP1^KD^ EVs) (**Sup. Fig. 4C-E**) and treating recipient cells with these. As expected, modulation of EFEMP1 levels in cells led to altered EFEMP1 levels in EVs isolated from these cells (**Sup. Fig. 4C-E**) and we found that EFEMP^OEX^ EVs promoted, and EFEMP1^KD^ EVs impaired migration and invasion of recipient BT549 cells (**Fig. 3A-E**). This was also true for MDA-MB-321 cells (**Sup. Fig. 4A, B**). To explore the contribution of EFEMP1 to the migration and invasion phenotypes induced by CIN^HIGH^ EVs, we induced CIN in control and EFEMP1^KD^ cells (**Sup. Fig. 4E**), isolated EVs, and determined their effects on recipient cell migration and invasion. While CIN^HIGH^ control EVs promoted migration and invasion as observed previously, this effect was largely abolished in CIN^HIGH^ EFEMP1^KD^ EVs (**Fig. 3F, G**), indicating that EFEMP1 plays a key role in the CIN^HIGH^ EV-mediated migration and invasion phenotypes observed in BT549 cells. We conclude that EFEMP1 directly promotes migration and invasion in TNBC cells, both independently and through EVs, and that the migration and invasion phenotypes driven by CIN^HIGH^ EVs are contingent upon EFEMP1 expression.

**Figure 3.**
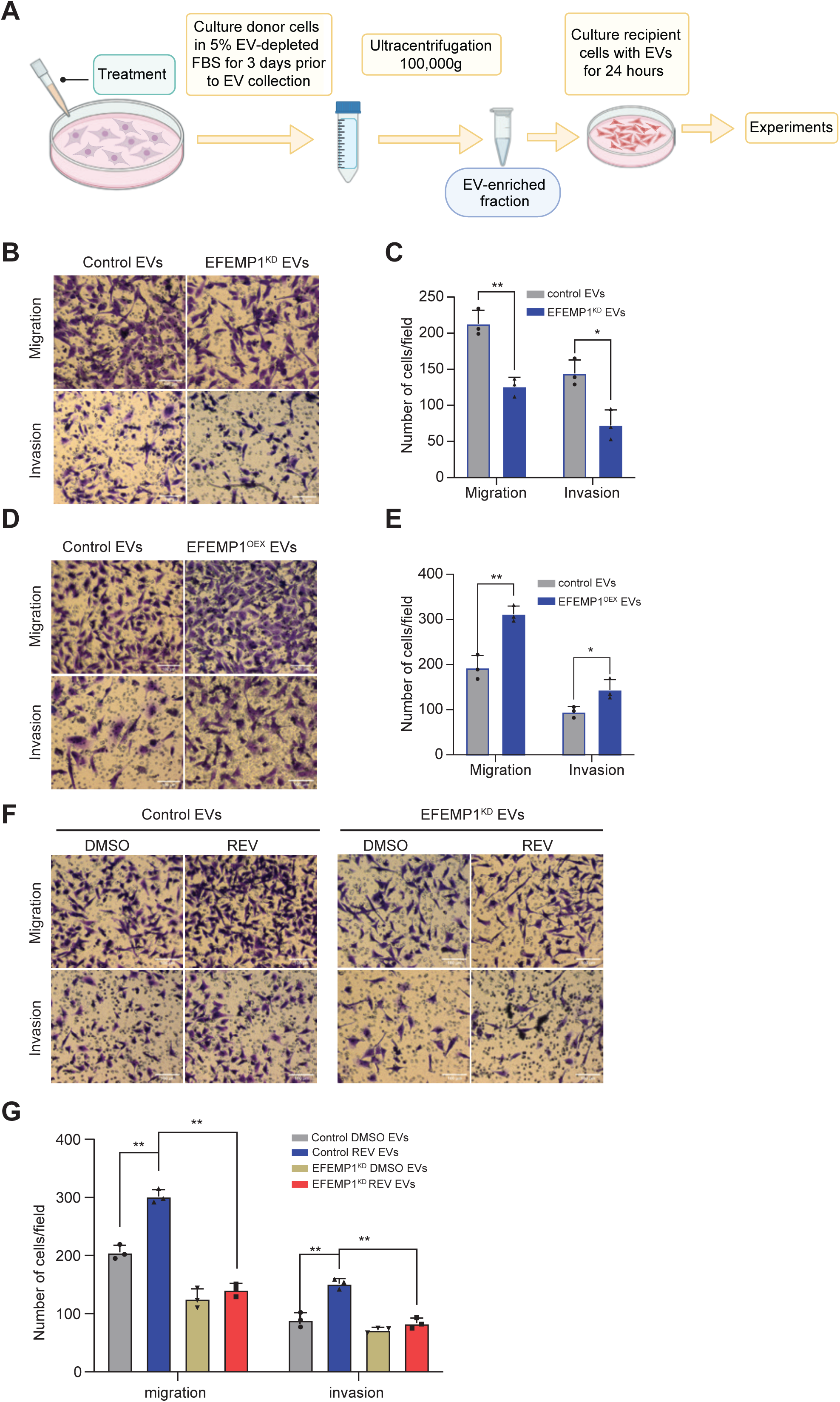
EFEMP1-enriched EVs promote cell migration and invasion. (**A**) Schematic outline of the treatment of BT549 cells with EVs and experimental setup for downstream analyses. (**B, C**) Representative images (**B**) and quantification (**C**) of BT549 migration and invasion Transwell experiments following EFEMP1 knockdown (EFEMP1^KD^) compared to control (scramble) EVs. Statistical significance was determined by a two-sided T-test (n= 3 biological replicates), *p < 0.05, **p < 0.01, ***p < 0.001, ****p < 0.0001. Scale bar: 100 µm. (**D, E**) Representative images (**D**) and quantification (**E**) of BT549 migration and invasion Transwell experiments following EFEMP1 overexpression (EFEMP1^OEX^) compared to control EVs. Statistical significance was determined by a two-sided T-test (n= 3 biological replicates), *p < 0.05, **p < 0.01, ***p < 0.001, ****p < 0.0001. Scale bar: 100 µm. **(F, G)** Representative images (**F**) and quantification (**G**) of migration and invasion of control and CIN^HIGH^ (REV-treated) BT549 cells using Transwell assays under Scramble control and EFEMP1^KD^ conditions. Statistical significance was determined by a two-sided T-test (n= 3 biological replicates), *p < 0.05, **p < 0.01, ***p < 0.001, ****p < 0.0001. Scale bar: 100 µm.

To better understand the mechanism underlying the invasion and migration phenotype instigated by EFEMP1 EVs, we performed RNA sequencing analysis on BT549 cells treated with control and EFEMP1^KD^ EVs. Among the 290 genes showing differential expression between cells treated with control EVs and EFEMP1^KD^ EVs, 29 have a known role in migration and/or invasion (**Sup. Fig. 5A**) (Shi *et al*, 2023). Notably, the dysregulated genes are enriched in pathways such as ’ECM receptor interaction’ and ’Focal adhesion’ (KEGG pathways, **Sup. Fig. 5B**) (Luo & Brouwer, 2013; Kanehisa *et al*, 2021; Ge *et al*, 2020). Specifically, genes involved in ’cell adhesion’ and ’cadherin binding’ (GO pathways, **Supp. Fig. 5C**) exhibit reduced expression levels in cells treated with EFEMP1^KD^ EVs compared to those treated with control EVs. Given the significant role of cell adhesion in cell migration (Khalili & Ahmad, 2015; Gupton & Waterman-Storer, 2006; Parsons *et al*, 2010), we next investigated the impact of EFEMP1^OEX^ and EFEMP1^KD^ EVs on the adhesion of BT549 cells. For this, wells were coated with diverse adhesive substrates, followed by addition of EV-treated BT549 cells. Cell adhesion was then assessed by inverting the plate and quantifying the adherent cells using crystal violet staining (**Sup. Fig. 5D**). These assays revealed that BT549 cells treated with EFEMP1^OEX^ EVs exhibited enhanced adhesion, while the converse was true for BT549 cells treated with EFEMP1^KD^ EVs (**Sup. Fig. 5E, F**). These findings suggest that EFEMP1-containing EVs promote adhesion of recipient cells, which could explain how EFEMP1 expression modulates migration and invasion behaviour induced by CIN^HIGH^ EVs (Khalili & Ahmad, 2015; Liu *et al*, 2019).

**Supplementary Figure 3.**
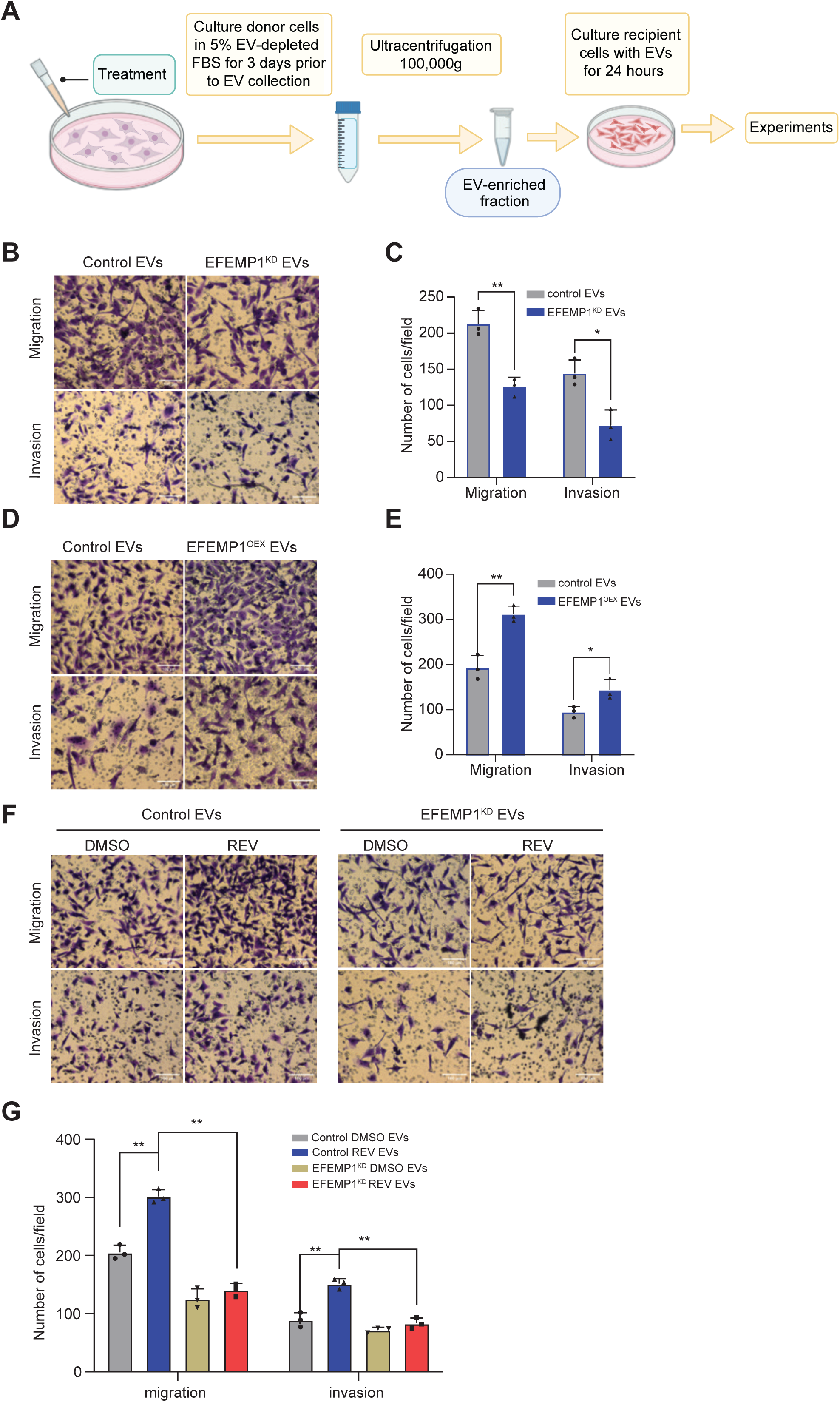
EFEMP1 enhances migration and invasion of BT549 and MDA-MB-231 cells. (**A**) qPCR quantification of EFEMP1 knockdown efficacy in BT549 cells: Statistical tests were done using a two-sided T-test (n=3; ***, p < 0.001). (**B**) qPCR quantification of EFEMP1 overexpression in BT549 cells. Statistical significance was assessed by a two-sided T-test (n=3; ****, p < 0.0001). (**C**) qPCR quantification of EFEMP1 knockdown efficiency in MDA-MB-231 cells. Statistical analysis using a two-sided T-test (n=3; ***, p < 0.001). (**D**) qPCR quantification of EFEMP1 overexpression. Statistical significance assessed by a two-sided T-test (n=3; ****, p < 0.0001). (**E-F**) Representative images (**E**) and quantification (**F**) of cell migration and invasion of BT549 cells with EFEMP1 overexpression assessed by trans-well assays. Statistical significance was determined by a two-sided T-test (n= 3 biological replicates), *p < 0.05, **p < 0.01, ***p < 0.001, ****p < 0.0001. Scale bar: 100 µm. (**G, H**) Representative images (**G**) and quantification (**H**) of cell migration and invasion of MDA-MB-231 cells with EFEMP1 overexpression assessed by trans-well assays. Statistical significance was determined by a two-sided T-test (n= 3 biological replicates), *p < 0.05, **p < 0.01, ***p < 0.001, ****p < 0.0001. Scale bar: 100 µm. (**I**) Representative images (left panel) and quantification (right panel) of scratch assays to quantify migration of BT549 cells with EFEMP1 knockdown. Statistical significance was determined by a two-sided T-test (n= 3 biological replicates), *p < 0.05, **p < 0.01, ***p < 0.001, ****p < 0.0001. (**J**) Representative images (left panel) and quantification (right panel) of scratch assays to quantify migration of BT549 cells with EFEMP1 overexpression. Statistical significance was determined by a two-sided T-test (n= 3 biological replicates), *p < 0.05, **p < 0.01, ***p < 0.001, ****p < 0.0001. (**K-L**) Representative images (**K**) and quantification (**L**) of cell migration and invasion of BT549 cells with EFEMP1 knockdown assessed by trans-well assays. Statistical significance was determined by a two-sided T-test (n= 3 biological replicates), *p < 0.05, **p < 0.01, ***p < 0.001, ****p < 0.0001. Scale bar: 100 µm. (**M, N**) Representative images (**M**) and quantification (**N**) of cell migration and invasion of MDA-MB-231 cells with EFEMP1 knockdown (two shRNAs) assessed by trans-well assays. Statistical significance was determined by a two-sided T-test (n= 3 biological replicates), *p < 0.05, **p < 0.01, ***p < 0.001, ****p < 0.0001.

**Supplementary Figure 4.**
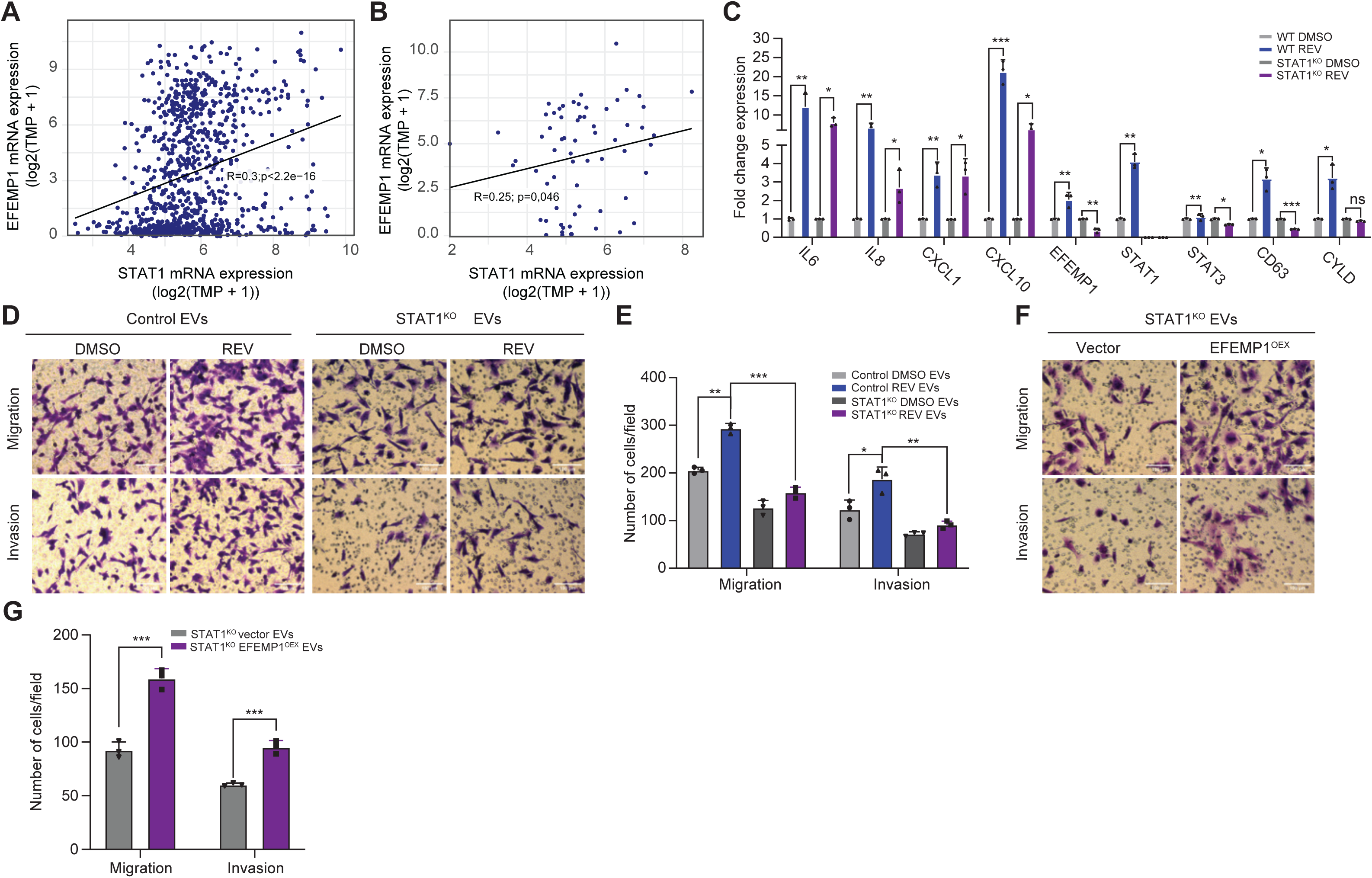
EFEMP1 levels in EVs modulate migration and invasion behaviour of MDA-MB-231 cells. (**A**) Representative images (left panel) and quantification (right panel) of cell migration and invasion of MDA-MB-231 cells treated with EFEMP1^KD^ EVs assessed by trans-well assays. Statistical significance was determined by a two-sided T-test (n= 3 biological replicates), *p < 0.05, **p < 0.01, ***p < 0.001, ****p < 0.0001. Scale bar: 100 µm. (**B**) Representative images (left panel) and quantification (right panel) of cell migration and invasion of MDA-MB-231 cells treated with EFEMP1^OEX^ EVs assessed by transwell assays. Statistical significance was determined by a two-sided T-test (n= 3 biological replicates), *p < 0.05, **p < 0.01, ***p < 0.001, ****p < 0.0001. Scale bar: 100 µm. (**C**) CD63, CD81, EFEMP1, Calnexin and beta-Actin protein levels in whole cell lysates and EVs from BT549 scramble and EFEMP1^KD^ cells determined by Western blot. (**D**) CD63, CD81, EFEMP1, Calnexin and beta-Actin protein levels in whole cell lysates and EVs from BT549 control and EFEMP1^OEX^ cells determined by Western blot. (**E**) CD63, CD81, EFEMP1, Calnexin and beta-Actin protein levels in whole cell lysates and EVs from DMSO- or REV-treated BT549 scramble and EFEMP1^KD^ cells determined by Western blot. (**F**) CD63, CD81, EFEMP1, Calnexin and beta-Actin protein levels in whole cell lysates and EVs from MDA-MB-231 scramble and EFEMP1^KD^ cells determined by Western blot. (**G**) CD63, CD81, EFEMP1, Calnexin and beta-Actin protein levels in whole cell lysates and EVs from MDA-MB-231 control and EFEMP1^OEX^ cells determined by Western blot.

**Supplementary Figure 5.**
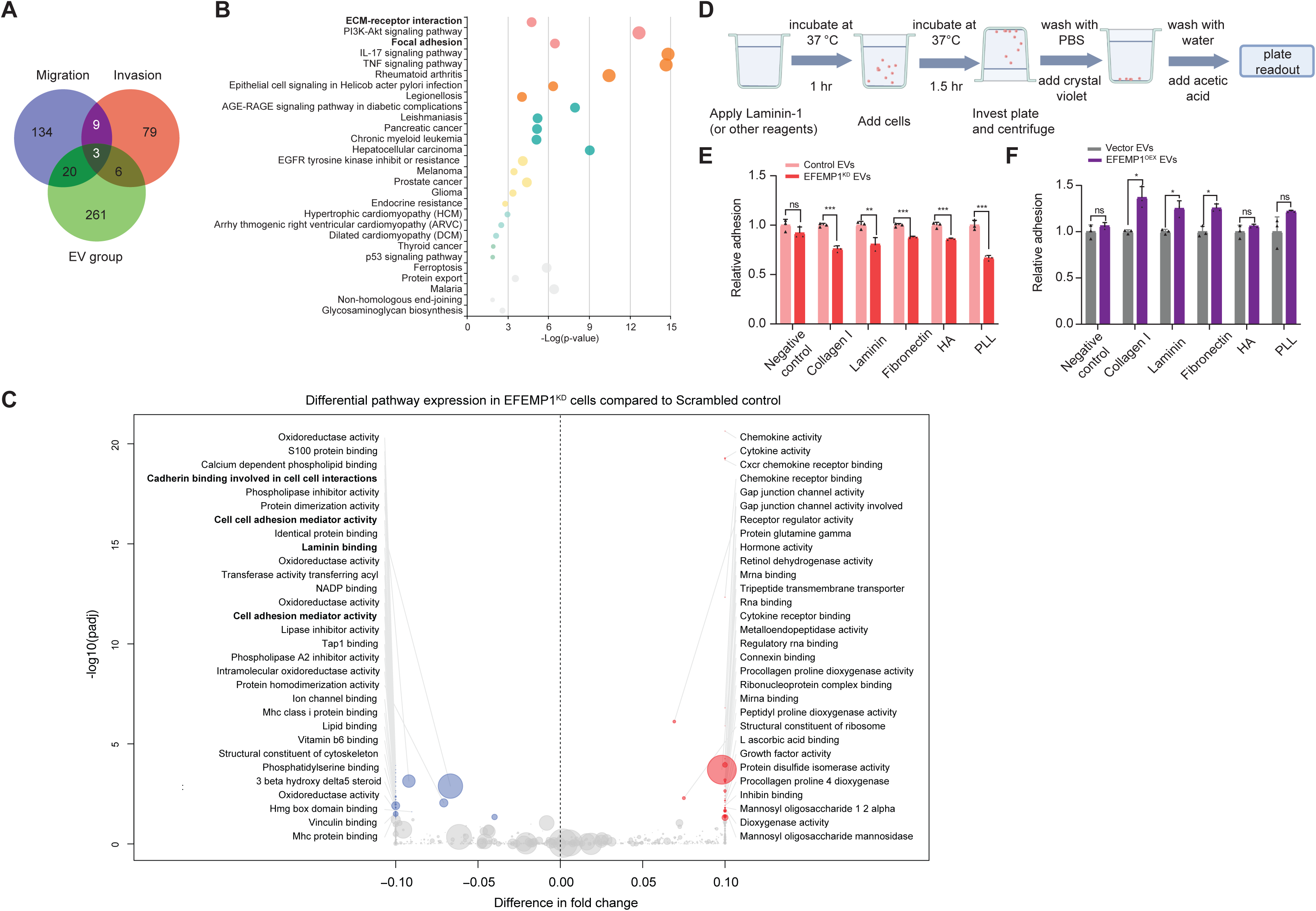
EFEMP1 modulates cell adhesion in a paracrine manner. (**A**) Venn diagram of RNA sequencing analysis illustrating the downregulation of 29 genes associated with migration and invasion in BT549 cells treated with EFEMP1^KD^ EVs. The gene sets for migration (166 genes) and invasion (97 genes) were derived from the Cancer Single-cell State Atlas, underscoring the significant role of EFEMP1 in regulating cellular migration and invasion patterns. (http://biocc.hrbmu.edu.cn/CancerSEA/goDownload). (**B**) KEGG pathway analysis of the 290 deregulated genes in EFEMP^KD^ cells. (**C**) Molecular Functions (MF) pathway analysis of the 290 deregulated genes in EFEMP^KD^ cells. (**D)** Schematic outline of the cell adhesion assay. (**E**) Quantification of cell adhesion of BT549 cells treated with EVs-derived from EFEMP1^KD^ cells. (**F**) Quantification of cell adhesion of BT549 cells treated with EVs-derived from EFEMP1^OEX^ cells.

### STAT1 is a regulator of EFEMP1 expression in CIN^HIGH^ cells

We recently identified STAT1 is a central node in CIN-induced inflammatory signalling (Schubert *et al*, 2021; Hong *et al*, 2022) and EFEMP1 was previously identified as a STAT1-target gene in a ChIP-seq experiment (Satoh & Tabunoki, 2013). Indeed, plotting EFEMP1 and STAT1 expression levels for > 1,000 cancer cell lines using DepMap data (dataset 23Q4; (Tsherniak *et al*, 2017)) confirmed a positive correlation between the expression of both genes across all DepMap included cancer cell lines (**Fig. 4A**), as well as all DepMap included breast cancer cell lines (**Fig. 4B**). To functionally test a possible epistatic relation between STAT1 and EFEMP1, we next generated STAT1^KO^ BT549 cells (**Sup. Fig. 6A**) and compared expression levels of several inflammatory genes (IL-6, IL-8, CXCL1, and CXCL10), and furthermore expression levels of EFEMP1, STAT1, STAT3, CD63 and CYLD in STAT1-proficient and -deficient backgrounds, with and without CIN. While cytokines were upregulated in a STAT1-independent manner, EFEMP1, STAT3, CD63 and CYLD all failed to be upregulated in STAT1-deficient BT549 cells, indicating that their expression is STAT1-regulated (**Fig. 4C**). In fact, EFEMP1, STAT3 and CD63 levels were even downregulated under CIN conditions. Also, note that STAT1-deficient cells show much lower basal expression of EFEMP1 (**Sup. Fig. 6A**). We then isolated EVs of STAT1-proficient and-deficient BT549 cells that were either treated with DMSO (CIN^LOW^) or MPS1 inhibitor (CIN^HIGH^), transferred these to recipient BT549 cells and quantified migration and invasion using trans-well assays. While CIN^HIGH^ EVs isolated from STAT1-proficient cells induced migration and invasion as expected, loss of STAT1 alleviated this phenotype, suggesting that STAT1-mediated EFEMP1 expression is required for this (**Fig. 4D, E**). To determine whether the decreased potential to induce migration and invasion of STAT1-deficient EVs was indeed dependent on EFEMP1, we overexpressed EFEMP1 in STAT1^KO^ BT549 cells, isolated EVs and compared migration and invasion between STAT1^KO^ cells with and without EFEMP1 overexpression. We found that re-expressing EFEMP1 in STAT1^KO^ BT549 was indeed sufficient to rescue the migration and invasion potential of EVs isolated from these cells (**Fig. 4F, G**). Jointly, these findings indicate that EFEMP1 is induced in a STAT1-dependent manner and that overexpression of EFEMP1 in STAT1^KO^ BT549 cells is sufficient for the secretion of EVs that promote migration and invasion of recipient TNBC cells.

**Figure 4.**
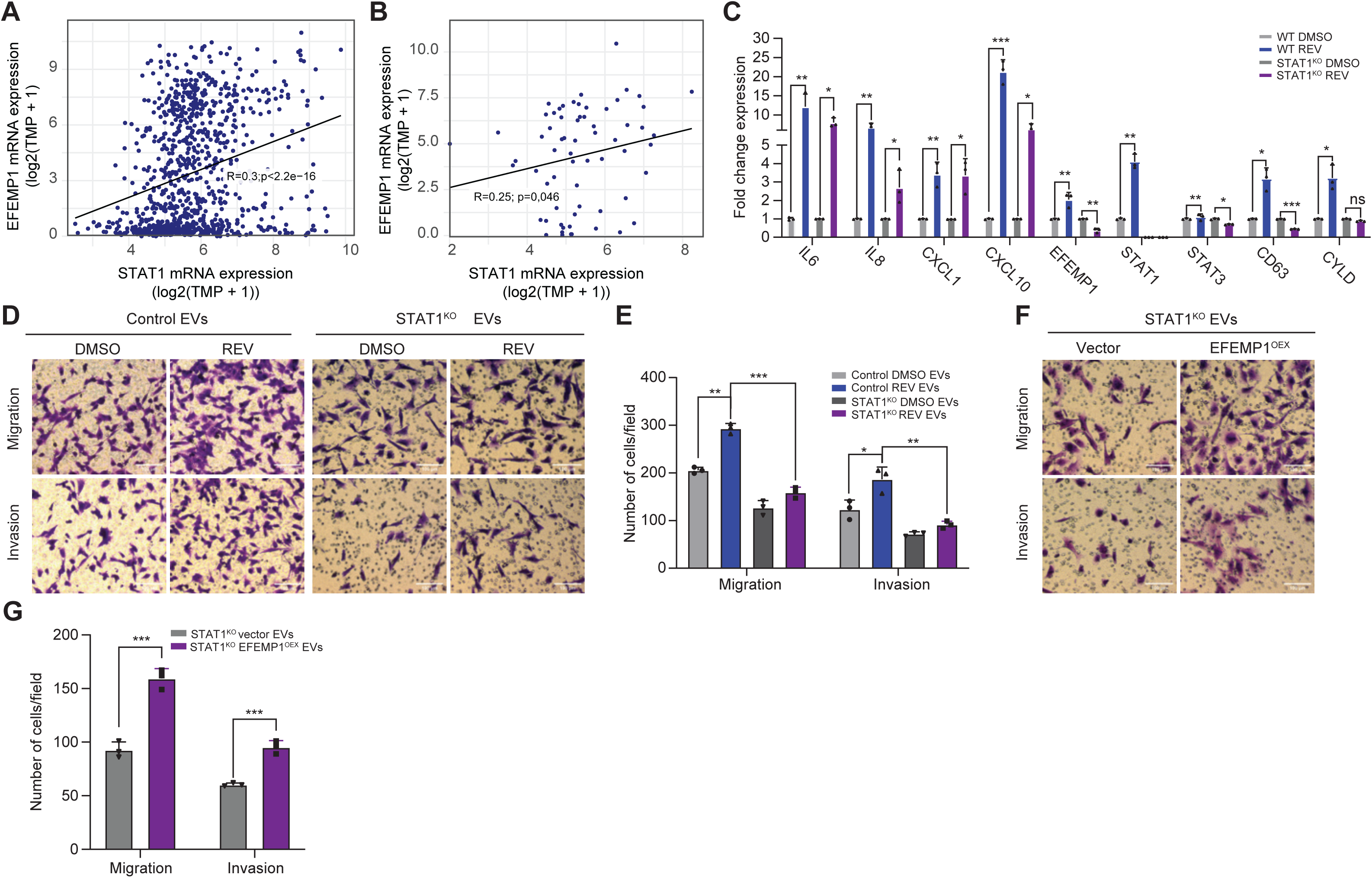
STAT1 is required for CIN-induced EFEMP1 expression and its downstream effects on migration and invasion. (**A**) Correlation analysis of EFEMP1 and STAT1 mRNA expression using DepMap data. (**B**) Correlation analysis of EFEMP1 and STAT1 mRNA expression in DepMap-included breast cancer cell lines. (**C**) qPCR quantification of IL6, IL8, CXCL1, CXCL10, EFEMP1, STAT1, STAT3, CD63 and CYLD in BT549 WT and STAT1 KO cells. Statistical analysis was done using two-sided T-test (N = 3; *p< 0.05, **, p< 0.01, ***p < 0.001, ****p < 0.0001). (**D, E**) Representative images (**D**) and quantification (**E**) of cell migration and invasion of BT549 cells co-cultured with EVs isolated from WT of STAT1^KO^ BT549 cells, treated with DMSO or REV, assessed by transwell assays. Statistical significance was determined by a two-sided T-test (n= 3 biological replicates), *p < 0.05, **p < 0.01, ***p < 0.001, ****p < 0.0001. Scale bar: 100 µm. (**F, G**) Representative images (**F**) and quantification (**G**) of cell migration and invasion of BT549 cells co-cultured with EVs isolated from STAT1^KO^ BT549 cells with or without EFEMP1 overexpression, assessed by transwell assays. Statistical significance was determined by a two-sided T-test (n= 3 biological replicates), *p < 0.05, **p < 0.01, ***p < 0.001, ****p < 0.0001. Scale bar: 100 µm.

**Supplementary Figure 6.**
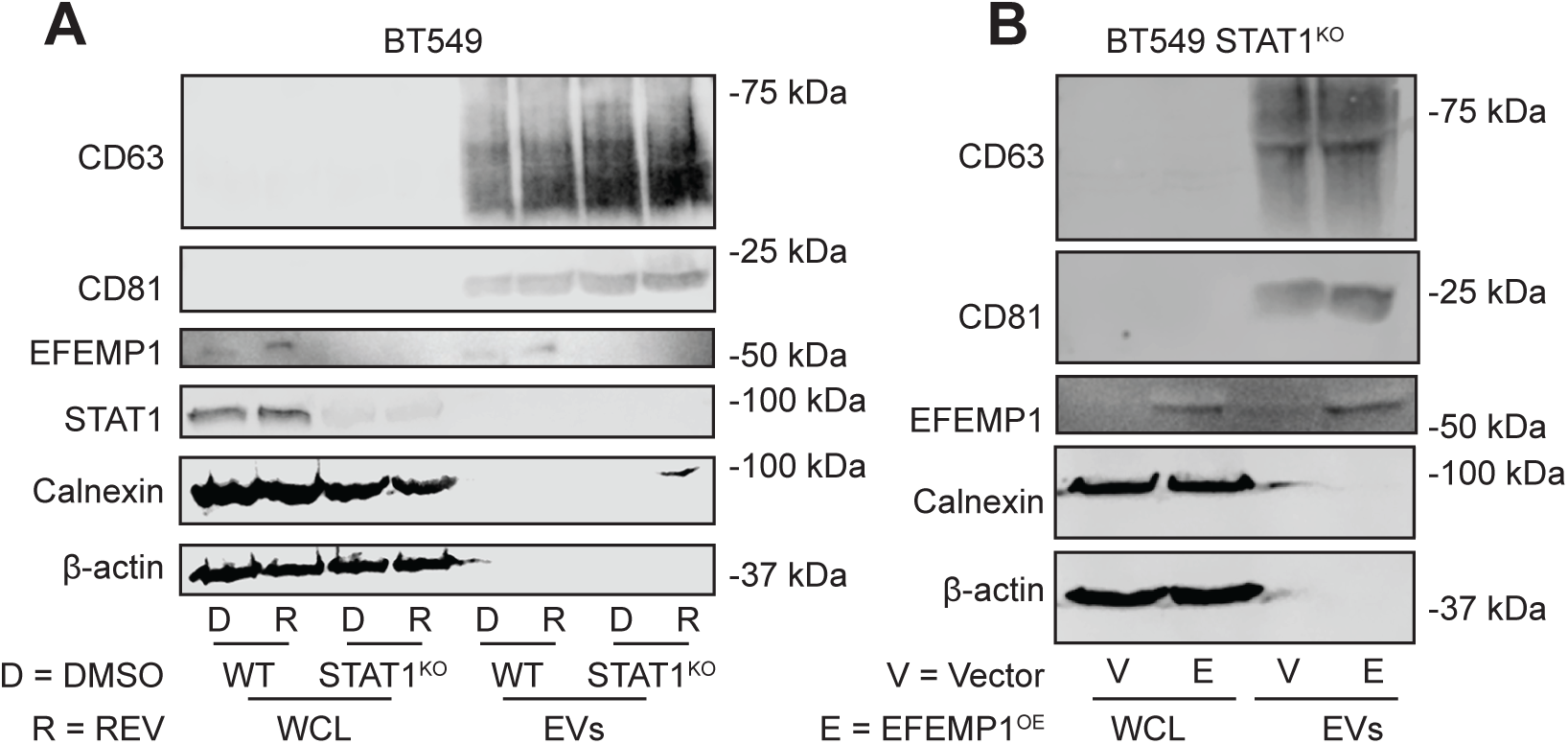
EFEMP1 and EV marker expression in BT549 WT and BT549 STAT1^KO^ cells. (**A**) CD63, CD81, EFEMP1, Calnexin and beta-Actin protein levels in whole cell lysates and EVs from BT549 and STAT^KO^ BT549 cells treated with DMSO or REV determined by Western blot. (**B**) CD63, CD81, EFEMP1, Calnexin and beta-Actin protein levels in whole cell lysates and EVs from STAT^KO^ BT549 cells treated with or without EFEMP1 overexpression determined by Western blot.

### EFEMP1-enriched EVs modulate tumour cell behaviour in a zebrafish xenograft model of MDA-MB-231 cells

Migration and invasion are important factors for metastasis (Pachmayr *et al*, 2017; Novikov *et al*, 2021; van Zijl *et al*, 2011).To test whether EFEMP1-containing EVs facilitate tumour cell spreading *in vivo*, we made use of a zebrafish xenograft model. For this, we cultured H2B-mCherry labelled MDA-MB-231 cells and treated them with EVs isolated from MDA-MB-231 cells in which EFEMP1 levels were modulated. Post-EV incubation, cells were injected into the perivitelline space (PVS) of 36 hpf-old zebrafish embryos (Rouhi *et al*, 2010; Teng *et al*, 2013; Martinez-Lopez *et al*, 2021). We then determined which fraction of the injected EV-treated MDA-MB-231 cells migrated to the tail section of the embryos (**Fig. 5A, Sup. Fig 7A-C**). We compared the spreading of cells treated with control EVs, EVs from EFEMP1^OEX^ and EVs from EFEMP1^KD^ cells (**Fig. 5B**) and found that while ∼50% of embryos injected with cells treated with control EVs showed cells spreading towards the tail of the embryo (**Fig. 5C**), this fraction was significantly reduced for cells treated with EFEMP1^KD^ EVs, and, conversely, significantly increased for cells treated with EFEMP1^OEX^ EVs. Quantifying the number of spreading cells further strengthened this observation: while ∼ 30% of the embryos injected with cells treated with control EVs showed more than 5 cells spreading to the tail, this fraction was reduced at least six-fold in embryos injected with cells treated with EFEMP^KD^ EVs (**Fig. 5D**). For cells treated with EFEMP1^OEX^ EVs, the proportion of embryos exhibiting the spread of more than 5 cells towards the tail doubled (**Fig. 5E**). Together, these data indicate that that EFEMP1-derived EVs can modulates cancer cell spreading *in vivo*.

**Figure 5.**
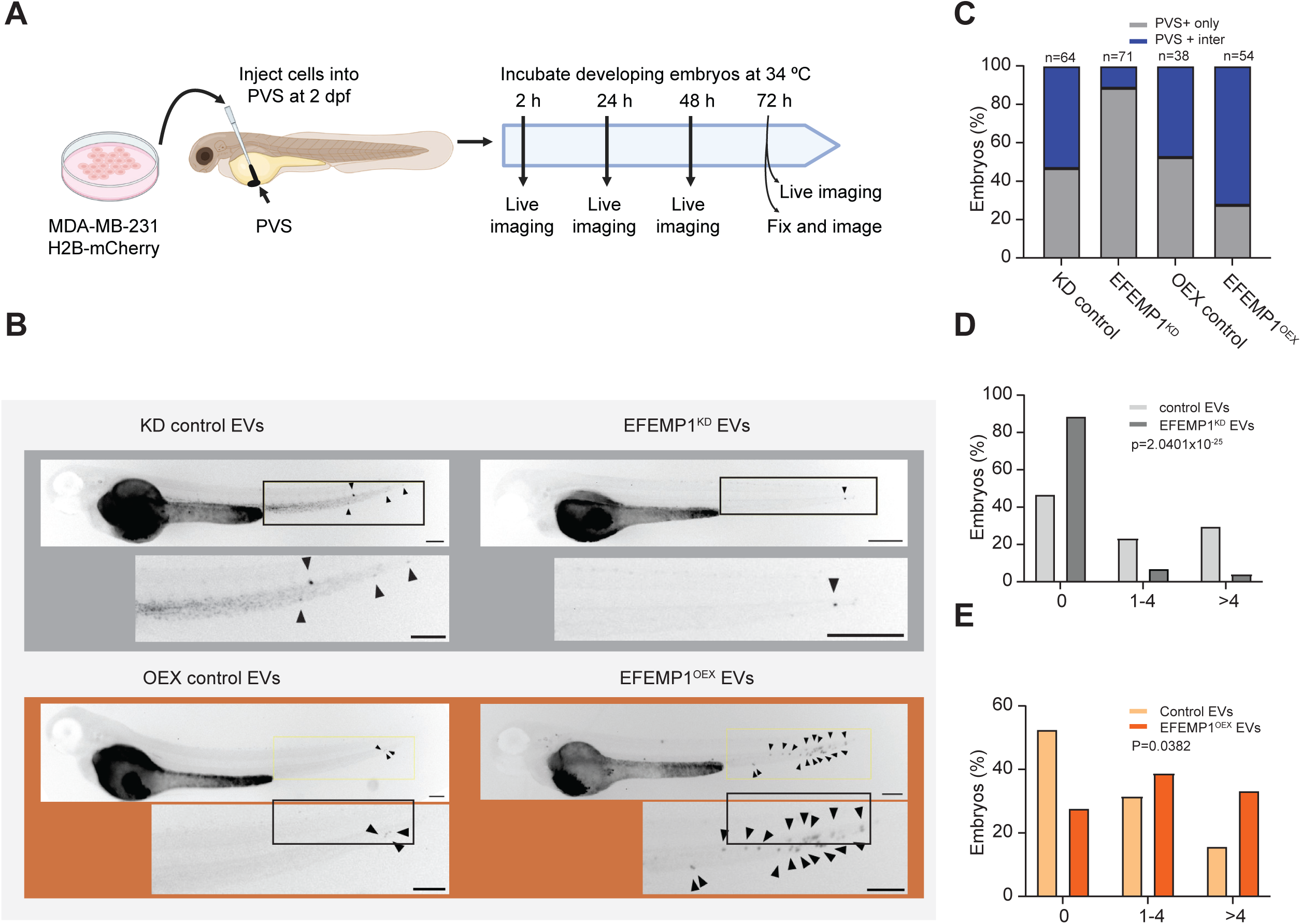
EMFEP1 promotes MDA-MB-231 cell spreading in a zebrafish xenograft model for metastasis. (**A**) Experimental setup for investigating the invasive behaviour of EV-treated MDA-MB-231 cells in zebrafish embryos. Xenografted embryos were monitored until 1-day post-transplantation (dpt). (**B**) Representative images of embryos xenografted with MDA-MB-231 cells treated with control (left column), EFEMP1^KD^ EVs (right column), or EFEMP1^OEX^ EVs (bottom rows) at 1 dpt. Arrowheads point to migrated MDA-MB-231 cells. Scale bar: 200 µm. (**C**) Quantification of MDA-MB-231 cell migration treated with EVs as indicated in large number of zebrafish embryos. Significance was tested using a chi-square test (****, p=3.4084×10^-30^). (**D**) Quantification of EV-treated MDA-MB-231 cell migration toward the zebrafish embryo’s tail region. Cells were treated with control or EFEMP1^KD^ EVs as indicated. Significance between groups (control or EFEMP^KD^ EVs) was tested using a chi-square test and migration patterns (categorized as 0, 1-4, and >4 cells migrated cells). p=2.0401×10^-25^. (**E**) Quantification of EV-treated MDA-MB-231 cell migration toward the zebrafish embryo’s tail region. Cells were treated with control or EFEMP1^OEX^ EVs as indicated. Significance between groups (control or EFEMP^KD^ EVs) was tested using a chi-square test and migration patterns (categorized as 0, 1-4, and >4 cells migrated cells). p=0.0382.

**Supplementary Figure 7.**
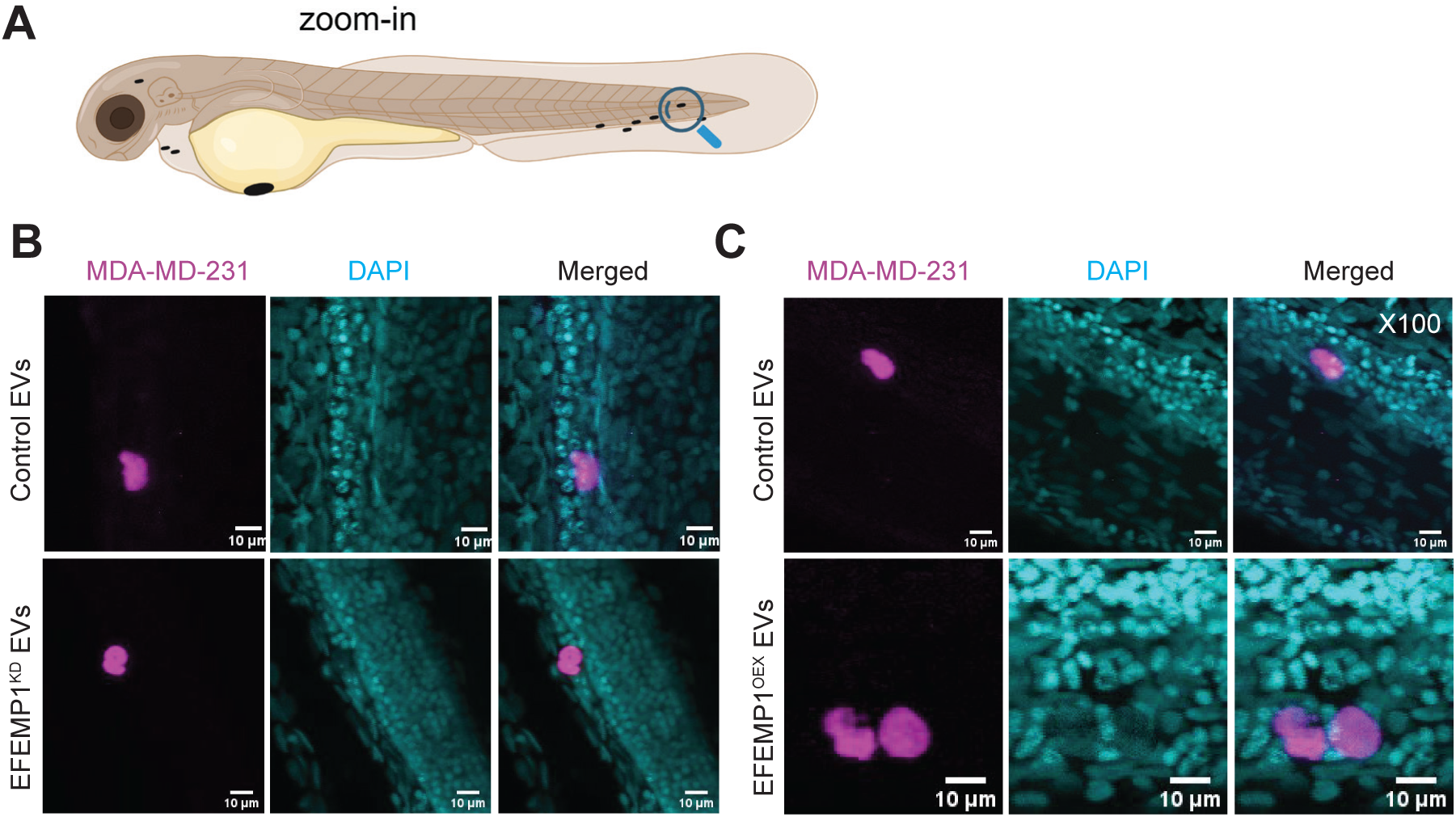
High resolution imaging confirms MDA-MB-231 spreading to the tail of xenografted zebrafish embryos. (**A**) Schematic of zebrafish embryo showing the region where migrated MDA-MB-231 cells were monitored. (**B, C**) Representative single optical section images of zebrafish embryo tails with mCherry-H2B labelled MDA-MB-231 cells 1 dpt. All nuclei were labelled with DAPI Scale bar: 10 µm.

### EFEMP1 and STAT1 expression patterns in breast cancer progression and prognosis

Finally, we assessed the relevance of our findings to human cancer. Using DepMap data (23Q4; (Tsherniak *et al*, 2017)), we determined the correlation between EFEMP1 mRNA levels and the aneuploidy score across all DepMap included cancer cell lines and across all DepMap included breast cancer cell lines and found a strong positive correlation for both (**Fig. 6A, B**). This is well in line with our experimental findings that CIN promotes EFEMP1 expression. Furthermore, when assessing TCGA patient survival we found that high expression of EFEMP1 is associated with a poorer survival for breast cancer across tumour grades (**Fig. 6C-F**), in agreement with our findings that high EFEMP1 expression promotes invasion and migration. We conclude that high expression of EFEMP1 is a poor prognosis marker in breast cancer, particularly from grade II and grade III breast cancer.

**Figure 6.**
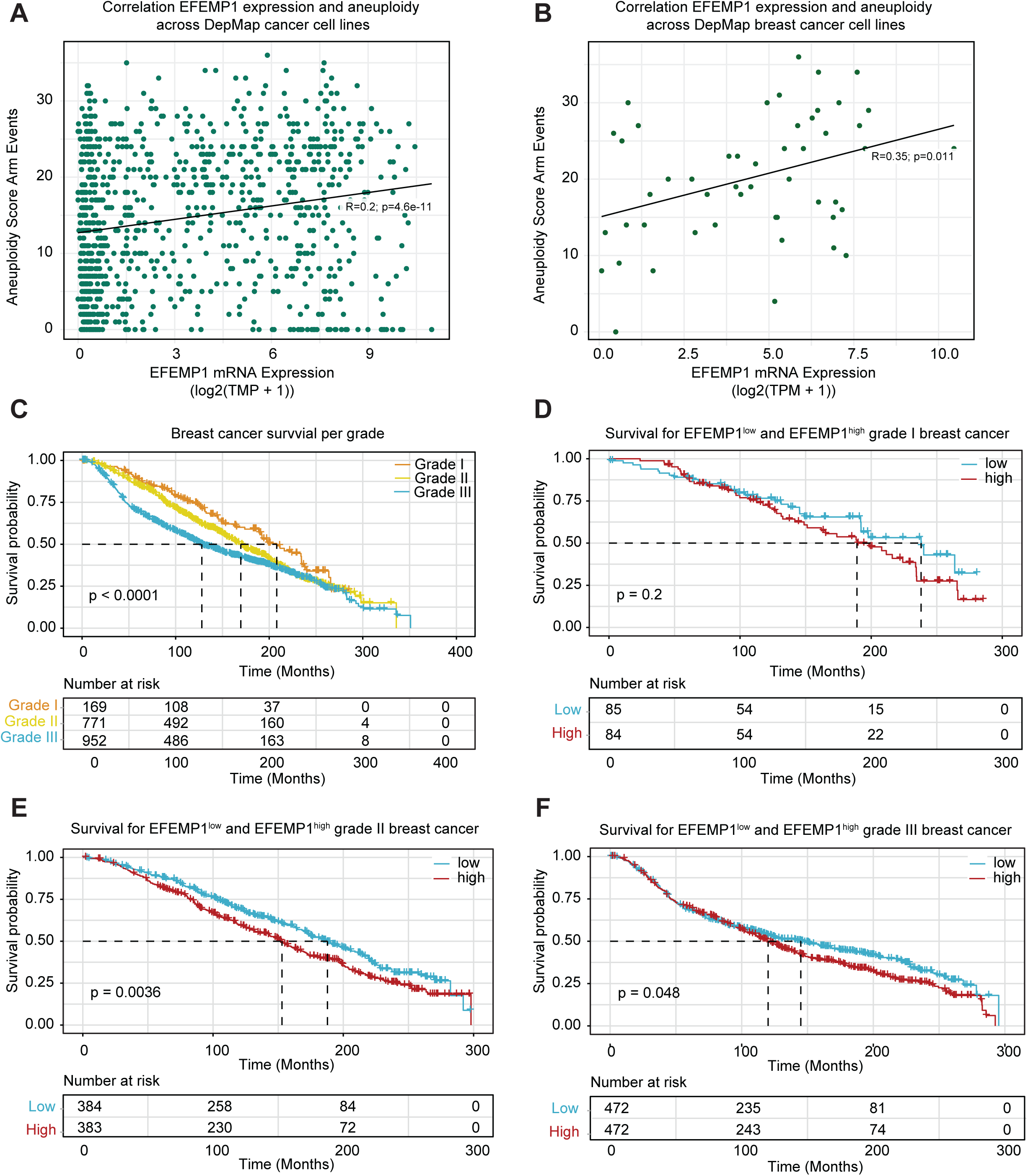
EFEMP1 and STAT1 expression patterns in breast cancer progression and prognosis. (**A-B**) Scatter plots showing the correlation between EFEMP1 expression and aneuploidy score in DepMap included cancer cell lines (**A**) or DepMap included breast cancer cell lines (**B**). (**C**) Kaplan-Meier survival curves for TGCA-included breast cancer patients, showing overall survival for grade 3 (blue) (*n* = 952), grade 2 (yellow) (*n=771*) and grade 1 (tangerine) (*n=169*) breast cancer. (**D-F**) Kaplan-Meier survival curves for TGCA-included breast cancer patients stratified for low or high EFEMP1 expression per grade (grade I, **D**; grade II, **E**, grade III, **F**). Significant differences between EFEMP1 expression groups were tested using a Log-rank Test.

While further work is required to explore the role of EFEMP1 as a cancer therapy target, altogether our work identifies EFEMP1 as a CIN-induced, STAT1-regulated factor that promotes migration and invasion of cancer cells in a paracrine manner. As high EFEMP1 expression is correlated with poor survival of breast cancer patients, future work should explore whether inhibition of EFEMP1 indeed improves breast cancer survival, for instance by reducing metastatic potential.

## Discussion

In this study, we investigated the paracrine effects of chromosomal instability using TNBC, the most aggressive breast cancer subtype, as a model cancer. We find that CIN, induced by the MPS1 inhibitor reversine promotes the secretion of extracellular vesicles, thus revealing a new mechanism by which CIN^HIGH^ cancer cells can influence the tumour microenvironment (TME). In line with our findings, others have reported that centrosome amplification, another driver of CIN, also leads to increased release of EVs (Adams *et al*, 2021).

We identify EFEMP1 as a key factor secreted by CIN^HIGH^ cancer cells via EVs to modulate the migration and invasion of recipient cells. EFEMP1 has previously been identified as a factor secreted by TNBC cells (McHenry & Prosperi, 2023), and as a biomarker of cancer detected in peripheral blood (Noonan *et al*, 2018). Our findings reveal EFEMP1 as a CIN-induced factor and a paracrine modulator of cell migration and invasion. While we show that EFEMP1-loaded EVs promote the migration of cancer cells in tissue culture and a zebrafish model, further work in for instance mouse models and human co-culture models is required to better understand which cell types in the TME are influenced by these EVs in a more human relevant setting and how this compares to the effects that the EVs have on neighbouring cancer cells. However, our finding that CIN-induced expression of EFEMP1 might contribute to metastasis aligns well with the discovery that CIN-driven activation of the cGAS–STING pathway promotes a pro-metastatic TME (Li *et al*, 2021a, 2023; Bakhoum & Cantley, 2018). Our work thus complements this recent work on the role of CIN in metastasis.

In addition to better understanding how CIN phenotypes influence the TME, our work further contributes to a broader understanding of how paracrine signalling of cancer cells can contribute to cell adhesion in cancer cell invasion and migration. Transcriptome analysis of TNBC cells treated with EVs isolated from EFEMP1^KD^ cells revealed a significant decrease in the expression of genes involved in adhesion, suggesting that EFEMP1 in EVs contributes to an increased adhesion phenotype in recipient cells, which was confirmed with adhesion assays of EMFEMP1^OEX^ and EFEMP1^KD^ cells. While an increased adhesion phenotype might appear at odds with the concomitant increased migration, this could be due to a dynamic regulation of cell-matrix interactions that enhances detachment and reattachment processes, thus offering a potential explanation for our finding that CIN promotes the migration of EV-recipient cells via EFEMP1. Indeed, prior work has identified a role for EFEMP1 in TNBC cell migration through its interaction with KISS1R (Noonan *et al*, 2018), albeit not yet in the context of CIN and EVs. Further work is required to better understand the molecular mechanism that underlies the CIN-induced EFEMP1-mediated effect on migration and invasion of cells.

We furthermore show that EFEMP1 expression is regulated by STAT1, a key factor in breast cancer progression (Chan *et al*, 2012), thus revealing a new regulatory pathway that regulates migration and invasion, downstream of CIN. Indeed, STAT1 was previously identified as a key signalling node downstream of CIN that is activated by acute CIN (Schubert *et al*, 2021; Hong *et al*, 2022), but also inactivated in CIN^HIGH^ cancers, presumably to prevent immune recognition of CIN^HIGH^ cancer cells. As CIN is known to promote metastasis, but also leads to downregulation of STAT1 signalling in cancer, it will be interesting to test whether EFEMP1 is upregulated in a STAT1-independent manner in CIN^HIGH^ cancers and thus contributes to the migration and invasion of cancer cells, and, via EVs, also to other cell types in the TME.

Our findings are supported by analyses of publicly available real-life datasets. Using DepMap data, we found that EFEMP1 expression positively correlates with aneuploidy in cancer cell lines. Furthermore, in line with our finding that CIN promotes EFEMP1 expression and that EFEMP1 expression promotes invasion and migration, analyses of TGCA data show that increased expression of EFEMP1 is associated with poor breast cancer patient survival.

Altogether, our work reveals EFEMP1 as a CIN-regulated factor that is secreted via EVs and that promotes cancer cell migration in a paracrine manner. Therefore, EFEMP1 might present a new clinical target of CIN^HIGH^ cancers to suppress increased metastasis rates observed for CIN^HIGH^ cancers.

## Materials and Methods

### Cell Culture

BT549 cells were cultured in RPMI 1640 (Gibco, 11554526) supplemented with 10% fetal bovine serum (FBS) (Thermo Fisher Scientific, 11573397) and 1% penicillin/streptomycin (P/S) (Gibco, 15140-122). MDA-MB-231 and 293FT cells were cultured in Dulbecco’s Modified Eagle Medium (DMEM) (Gibco, 31966-021) with 10% FBS and 1% P/S. All cell lines were obtained from the American Type Culture Collection (ATCC). Cell lines were grown at 37°C and 5% CO_2_. For passaging, cells were detached using Tryple Express (Gibco, 12605-010).

### Immunofluorescence Microscopy

For immunofluorescence assays, cells were seeded on glass coverslips in 24-well plates. Following 24 hours of treatment, cells were fixed with 4% formaldehyde (Sigma-Aldrich, 252549) for 15 minutes and washed twice with cold PBS (Gibco). Permeabilization was performed using 0.1% Triton X-100 (Sigma-Aldrich, 9036-19-5) in PBS for 5-10 minutes at room temperature. After blocking with 3% FBS in PBS for 30 minutes, cells were incubated overnight with a CD63 antibody (H5C6; Novus Biologicals, NBP2-42225) diluted in the blocking buffer. Thereafter, cells were washed and incubated with a secondary anti-mouse Alexa Fluor 488 antibody (Invitrogen, A-21110) and Phalloidin-iFluor 555 (Abcam, ab176756) for F-actin staining. Nuclei were stained with DAPI (Sigma, D9542-1MG). Imaging was conducted using a Leica Sp8 confocal microscope. CD63 intensity was quantified using ImageJ (v1.53).

### EV Isolation

EVs were purified from the culture medium, supplemented with 5% EV-depleted FBS, as previously described (Théry *et al*, 2018). Cells were cultured to reach 50-60% confluence before the medium was replaced with fresh medium containing EV-depleted FBS. Cells were then incubated for an additional 2-3 days. To isolate EVs, we employed a dual approach: centrifugation (Merchant *et al*, 2017; Théry *et al*, 2006) and the use of EVs isolation tubes (Vivaspin® 20, 100 kDa MWCO Polyethersulfone, Cytiva, 28-9323-63), allowing for a comprehensive and effective purification process.

### EdU Proliferation Assay

For EdU incorporation assays, cells were plated on coverslips in 24-well plates and treated with EVs for 24 hours. For S-phase labelling, cells were pulsed with 10 µM EdU (Lumiprobe,10540) for 2 hours, fixed in 4% formaldehyde (Sigma-Aldrich, 252549), and EdU was detected using click chemistry (Lumiprobe, D1330). Slides were imaged by fluorescence microscopy (Olympus IX51 or BX43), with image analysis and quantification performed using Fiji (ImageJ v1.53c).

### Western Blot Assays

Cells or extracellular vesicles (EVs) were lysed using RIPA buffer (150 mM Sodium chloride, 50 mM Tris-HCl at pH 8.0, 1% NP-40, 0.5% Sodium deoxycholate, and 0.1% SDS), complemented with a protease inhibitor cocktail (Roche) for 30 minutes. The cell lysates were combined with 5X SDS buffer, while 4X LDS buffer was utilized for EV lysates. Samples were heated at 95°C for 5 minutes to denature proteins.

Protein concentration was quantified using the BCA assay (Pierce™ BCA Protein Assay Kits, 23225, Thermo Scientific™). For this, 20-50 µg of protein was loaded per sample onto 7.5-10% polyacrylamide gels for electrophoretic separation. Following electrophoresis, proteins were transferred onto PVDF membranes, which were then blocked using Odyssey blocking buffer (LI-COR Biosciences) or 5% non-fat milk in TBS-0.1%T to minimize non-specific binding.

The primary antibodies for Western blots used in this study were EFEMP1 (1:300, Novus Biologicals, NBP1-77040), β-Actin (1:2000, Cell Signaling Technology, 4970 or 3700S), STAT1 (1:1000, Cell Signaling Technology, 9172S), CD63 (1:1000, Novus Biologicals, NBP2-42225), CD81 (1:1000, Novus Biologicals, NB100-65805SS), and Calnexin (1:1000, Novus Biologicals, NB100-1965SS). All antibodies were incubated with the blots overnight at 4°C.

Secondary antibodies used were IRDye 800CW Goat anti-Rabbit IgG (H+L) (1:15000, LI-COR) and IRDye 680CW Goat anti-Mouse IgG (H+L) (1:15000, LI-COR), or for HRP-conjugated antibodies, Amersham ECL Mouse IgG, HRP-linked whole Ab (1:5000, Cytiva, NA931) and Amersham ECL Rabbit IgG, HRP-linked whole Ab (1:5000, Cytiva, NA934). These were incubated for 1 hour at room temperature.

Blots incubated with IRDye-conjugated antibodies were imaged using the Odyssey imaging system and analysed using Image Studio Lite software (LI-COR Biosciences). For HRP-conjugated antibodies, protein signals were detected using the ECL Chemiluminescent Kit (Thermo Fisher Scientific) and imaged with an Amersham Imager 600 (GE Healthcare).

### Trans-well assays

To assess cell migration, cells were seeded directly onto transwell chambers (cellqart, 9328002). For invasion assays, ECM gel (Corning, 356234) was prepared and applied to cool Millicell inserts (cellqart, 9328002) to solidify at 37 °C for two hours (Vasudevan *et al*, 2020). After a PBS wash, cells suspended in serum-free RPMI were seeded into the upper compartment of both setups. The lower compartment contained RPMI (or DMEM) with 10% FBS as a chemoattractant. Post-incubation, cells on the lower membrane side were fixed, stained with crystal violet, and imaged. Quantitative analysis involved averaging cell counts from three randomly selected fields per membrane at 100x magnification, using an Olympus IX51 fluorescence microscope. Each assay was replicated in triplicate.

### Scratch Assays

For scratch assays, cells were detached, counted, and seeded in 6cm dishes. For gene knockdown, cells were treated with doxycycline, while gene overexpression was achieved via plasmid transfection, both for 2-3 days. Cells (3.5×10^5^ for BT549) were then reseeded in 12- well plates. After cell attachment, a 1mm pipette tip was used to create a straight scratch. Post-scratching, the monolayer was washed with PBS and replenished with an FBS-free medium. Scratch closure was monitored using an IncuCyte machine at 10x magnification, capturing images every 2 hours for 24-48 hours. Image analysis and quantification were performed using Fiji software (ImageJ 1.53C).

### Zebrafish Xenograft Experiments

Zebrafish xenograft experiments were performed according to the European animal welfare regulations and standard protocols. Transparent Casper fish embryos were housed at 31°C until transplantation. To xenograft MDA-MB-231 cells, an established perivitelline space (PVS) xenograft method was used (Rouhi *et al*, 2010; Teng *et al*, 2013; Martinez-Lopez *et al*, 2021). Prior to xenografting, MDA-MB-231 WT cells expressing H2B-mCherry were pre-treated with EVs for 3 days, and then harvested for microinjection. At 36 hpf, healthy embryos were dechorionated, anaesthetized with tricaine, and arranged in agarose dishes. Cells suspended at 0.12×10^6^ cells/μL were loaded into capillaries with filament and microinjected into the PVS using a pneumatic pico pump (PV820, World Precision Instruments). This process was completed within 1.5 hours.

Post-xenograft embryos were transferred to clean E3 medium dishes and examined within 2 hours. Criteria for inclusion in the study included embryos with cells solely in the PVS. Embryos were incubated at 34°C in E3 medium in individual wells of a 24-well plate for imaging until fixation.

To assess tumour cell dissemination, live embryo images were captured at 2- and 24-hours post-transplantation using either a Leica MZ FLIII fluorescence stereomicroscope with a Leica DFC3000G digital camera or a Zeiss Axio Zoom V16 fluorescence stereomicroscope with an HRm digital camera. For higher resolution imaging, fixed xenograft embryos were processed by permeabilization using Proteinase K (10 μg/ml in PBS, Lot: 30032017, Roche) for 15-45 minutes, depending on embryo age, at 37°C. Subsequently, the embryos were stained with DAPI (Sigma, D9542-1MG), mounted, and subjected to imaging. The tail regions of embryos harbouring migrated cells were imaged using a Leica SP8x confocal microscope equipped with a 40X oil objective.

### Transmission Electron Microscopy (TEM)

For TEM, isolated EVs were fixed using 2% paraformaldehyde in 0.1 M sodium cacodylate buffer at pH 7.4 (Théry *et al*, 2018; Joshi *et al*, 2020). Grids were prepared with a double layer of Formvar film and carbon coating for enhanced stability. These grids were treated with 1% Alcian blue in 1% acetic acid, followed by thorough rinsing with double-distilled water. 10 µL of the EV suspension was applied to the hydrophilic grid surface and allowed to adhere for 30 minutes. After washing, the grids were rapidly stained with either 0.5% uranyl acetate or 1% phosphotungstic acid at a neutral pH, crucial for enhancing the EVs’ contrast. The samples were incubated at room temperature for 20 minutes to allow for full interaction between the stain and the EVs and then examined under a Transmission Electron Microscope (Talos200 TFS) (Théry *et al*, 2018; Joshi *et al*, 2020).

### Nanoparticle Tracking Analysis (NTA)

The size and concentration of the EVs were determined using a PMX-130 MONO Laser ZetaView® (Particle Metrix, Ammersee, Germany), with data analysis conducted via ZetaView 8.04.02 software. Calibration was performed using 110 nm polystyrene particles, and measurements were kept consistent at 25°C. EV samples were diluted in 1×PBS buffer for analysis. Nanoparticle tracking measurements were recorded and analysed at 11 distinct locations per sample. Each experiment was repeated three times.

### CRISPR-Cas9 Mediated Gene Knockout

To create CRISPR knockout cell lines targeting the human STAT1 gene (details in Supplementary Table 1), we utilized the pSpCas9(BB)-2A-Puro V2.0 (PX459) plasmid (Addgene plasmid 62988, Feng Zhang) (Ran *et al*, 2013). sgRNAs were cloned into this Cas9 plasmid using BbsI (NEB). STAT1 knockouts were made by transfecting with these sgRNA plasmids using FuGENE® HD Transfection Reagent (Promega, E2311). Cells underwent puromycin selection (Invitrogen, 0.1-1µg/ml) for 5 days to select for for knockouts. gRNA sequences are provided in Supplementary table2.

### shRNA Mediated Knockdown

For shRNA experiments, we used a Tet-pLKO-puro vector (Addgene plasmid 21915) (Wiederschain *et al*, 2009). shRNAs were cloned using AgeI and EcoRI restriction enzymes (NEB) and verified by sequencing. Target cells were transduced with these lentiviral constructs and selected using puromycin for 3-7 days. shRNA expression was induced with 1 µg/mL doxycycline (Sigma Aldrich) in the culture medium. Gene knockdown efficacy was confirmed through quantitative RT-PCR after three days of induction, and protein level changes were assessed by western blotting. The shRNA sequences are provided in Supplementary Table2.

### Protein overexpression

For labeling and tracking the uptake of extracellular vesicles (EVs), we utilized the pLenti-pHluorin_M153R-CD63-mScarlet plasmid (Addgene, Plasmid 172118). This dual-color fluorescent reporter is specifically designed for identifying CD63-positive exosome secretion and uptake. Our procedures adhered closely to the protocols provided by Addgene for plasmid 172118 (Sung *et al*, 2020). EFEMP1 overexpression was achieved by transduced cells with FL-fibulin-3 pcDNA4 (Addgene, Plasmid 29700) (Hu *et al*, 2009) plasmid using FuGENE® HD Transfection Reagent (Promega, E2311).

### Lentiviral Transduction

Lentiviruses were produced by transfecting 293FT cells with the selected vector and essential packaging plasmids (pSPAX2 and pMD2.G, gifts from Didier Trono; Addgene plasmid 12260 and 12259). Post-transfection, the medium was harvested, filtered through a 0.45 µm filter, and used for transducing target cells in the presence of 4-8 µg/ml polybrene (Sigma-Aldrich).

### Extraction of RNA and qRT-PCR

For qPCR analysis, RNA was extracted using the RNA plus isolation kit (Qiagen, 74136). Primer sequences are listed in Supplementary Table 1. For cDNA synthesis, 1.5 µg of RNA was added to a LunaScript RT SuperMix Kit (Bioke) in a 20 µl reaction. qPCR was performed using iTaq Universal SYBR Green supermix (Biorad) on a LightCycler® 480 Instrument. Data analysis was conducted using Excel (Microsoft) and Prism software (GraphPad). The qPCR primers are provided in Supplementary Table 2.

### Time-lapse imaging

Chromosomal abnormalities were quantified using time-lapse imaging. For this, cell lines were transduced with lentiviral H2B-mCherry constructs. One day prior to imaging, BT549 or MDA-MB-231 cells were seeded into imaging disks (Greiner Bio-One, 627870). Imaging was performed on a DeltaVision Elite microscope (GE Healthcare), fitted with a CoolSNAP HQ2 camera and a 40X, 0.6 NA immersion objective lens (Olympus) for at least 20 hours with images captured every 6 minutes. Each imaging stack consisted of 30 to 40 Z-stacks at 0.5 µm apart. Images were analysed using ICY software (Institut Pasteur). Only mitotic cells were included in the analyses. For live-cell imaging, the cells were pretreated with the drug one hour before initiating the imaging sessions.

### Scoring System for Chromosomal Instability

To streamline the quantification of CIN phenotypes in your method, we categorized mitotic events based on severity and complexity, following the standards set by previous studies (Crozier *et al*, 2022; Thu *et al*, 2018; Huis In ’t Veld *et al*, 2019), Chromosomal events were scored on a scale from 0 to 5, with ’other’ events representing atypical aberrations scored variably (1-5) but excluded from main statistical analysis due to their unique nature. In Figure 1A-C, these events are colour-coded:

0 points (green): Correct Chromosomal Segregation.

1 point (blue): DNA Bridge, Micronucleus Formation, Chromosomal Lagging.

2 points (yellow): DNA Bridge/Lagging with Micronucleus, Metaphase Misalignment.

3 points (orange): Metaphase Misalignment with Micronucleus or Lagging/Bridge.

4 points (purple): Metaphase Skipping, Mitotic Failure, Complex Misalignment.

5 points (red): Metaphase Skipping with DNA Bridging or Micronucleus Formation. Other (1-5 points, grey): Non-standard chromosomal aberrations or anomalies.

### Mass spectrometry

Coomassie-stained bands were excised in one gel slice that was further cut into small pieces and destained using 70% 50 mM NH_4_HCO_3_ and 30% acetonitrile. The reduction was performed using 10 mM DTT dissolved in 50 mM NH4HCO3 for 30 min at 55°C. Next, the samples were alkylated using 55 mM chloroacetamide in 50 mM NH_4_HCO_3_ for 30 min at room temperature and protected from light. Subsequently, samples were washed for 10 min with 50 mM NH_4_HCO_3_ and for 15 min with 100% acetonitrile. The remaining fluid was removed, and gel pieces were dried for 15 min at 55°C. Tryptic digestion was performed by the addition of sequencing-grade modified trypsin (Promega; 25 µL of 10 ng/mL in 50 mM NH_4_HCO_3_) and overnight incubation at 37°C. Peptides were extracted using 5% formic acid in water, followed by a second elution with 5% formic acid in 75% acetonitrile. Samples were dried in a SpeedVac centrifuge and dissolved in 20 µL 5% formic acid in water for analysis with LC-MS/MS.

The samples were analyzed on a nanoLC-MS/MS consisting of an Ultimate 3000 LC system (Thermo Fisher Scientific, USA) interfaced with a Q Exactive Plus mass spectrometer (Thermo Fisher Scientific). Peptide mixtures were loaded onto a 5 mm × 300 μm i.d. C18 PepMAP100 trapping column with water with 0.1% formic acid at 20 μL/min. After loading and washing for 3 min, peptides were eluted onto a 15 cm × 75 μm i.d. C18 PepMAP100 nanocolumn (Thermo Fisher Scientific). A mobile phase gradient at a flow rate of 300 μL/min and with a total run time 120 min was used: 2% − 45% of solvent B in 92 min; 45% − 85% B in 1 min; 85% B during 6 min, and back to 2% B in 0.1 min. Solvent A was 100:0 water/acetonitrile (v/v) with 0.1% formic acid, and solvent B was 0:100 water/acetonitrile (v/v) with 0.1% formic acid. In the nanospray source a stainless-steel emitter (Thermo Fisher Scientific) was used at a spray voltage of 2 kV with no sheath or auxiliary gas flow. The ion transfer tube temperature was 250 °C. Spectra were acquired in data-dependent mode with a survey scan at m/z 375 − 1600 at a resolution of 70 000 followed by MS/MS fragmentation of the top 10 precursor ions at a resolution of 17 500. Singly charged ions were excluded from MS/MS experiments, and fragmented precursor ions were dynamically excluded for 20 s.

PEAKS Studio version Xpro (Bioinformatics Solutions, Inc., Waterloo, Canada) software was used to search the MS data against the UniProt reviewed human protein sequence database (19-12-2023, 26450 entries) with the UniProt reviewed bovine protein database as contaminant database. Search parameters: trypsin digestion with up to 2 missed cleavages; fixed modification carbamidomethylation of cysteine; variable modification oxidation of methionine and deamidation of asparagine and glutamine; precursor mass tolerance of 15 ppm; fragment mass tolerance of 0.02 Da. The false discovery rate was set at 0.1% on the peptide level. Label free quantitation was performed with the PEAKS Q module, with the 3 reversine-treated cell samples and the 3 DMSO-treated cells assigned as separate groups, and with a significance threshold of 13.

### Analysis of Differentially Expressed Proteins

To determine differential proteins expression from mass spectrometry results, we utilized Morpheus (https://software.broadinstitute.org/morpheus) for heatmap generation. This analysis particularly highlighted the top 20 upregulated proteins in EVs derived from reversine-treated cells and revealed EFEMP1 as a significantly enriched protein (1.38-fold change, p=0.021). Furthermore, the cellular component enrichment of these differentially expressed proteins was analysed using Funrich 3.1.4.

We used ShinyGO v0.80 (http://bioinformatics.sdstate.edu/go/) for gene ontology analyses (Ge *et al*, 2020). These analyses were complemented by pathway clustering and protein– protein interaction network analyses using STAGEs (https://kuanrongchan-stages-stages-vpgh46.streamlit.app/).

### RNA Sequencing Sample and Library Preparation

RNA quality control (QC) involved concentration assessment using Nanodrop and quality and RIN value measurement via HSRNA screentape assay or RNA screentape assay on the Agilent 4200 Tapestation system.

Library preparation employed the Smart-3seq method on an Agilent Bravo Automated Liquid Handling Platform, using 100 ng input per sample. For lower-concentration samples, a pre-bead clean-up (poly A RNA capturing) with SPRI beads was performed. Indexes utilized were NEBNext Multiplex oligos for Illumina (E6440L) with 12 PCR cycles. Bead clean-up was done twice at a 0.8x ratio.

Post-library preparation QC included concentration measurement using Qubit 1X dsDNA HS Assay Kit and BioTek Synergy H1 Multimode Reader, with molarity assessment via HSD5000 or D5000 screentape assay on the Agilent 4200 Tapestation system.

Libraries were diluted to 4 nM and pooled. Superpool concentration and molarity verification employed the Qubit 1X dsDNA HS Assay Kit, Qubit 4 Fluorometer, and HSD5000 screentape assay.

Sequencing was conducted on the Illumina Nextseq2000 system using a P2 100 cycle kit. Run specifications included a read 1 of 113 bp, read 2 of 7 bp, index 1 of 9 bp, index 2 of 9 bp, loaded molarity of 750 pM, and a phiX percentage of 5%.

### RNA Sequencing Results Analysis

We analysed RNA sequencing data by comparing differentially expressed genes (>2-fold) in EFEMP^KD^ cells to on gene sets related to migration (166 genes) and invasion (97 genes), obtained from the Cancer Single-cell State Atlas (http://biocc.hrbmu.edu.cn/CancerSEA/goDownload) (Shi *et al*, 2023). The overlap among these gene sets was visualized using Venn diagrams visualized using an online tool (https://bioinformatics.psb.ugent.be/webtools/Venn/). For the GO gene set enrichment within the RNA sequencing results, we performed KEGG pathway analysis using KOBAS (http://bioinfo.org/kobas/genelist/) (Bu *et al*, 2021), focusing on pathways with P<0.05.

### Cell adhesion assays

For cell adhesion assays (Khalili & Ahmad, 2015), 96-well plates were first coated with human fibronectin (5 μg/mL; CellSystems; 5050), mouse laminin (5 μg/mL; Thermo Fisher; 23017015), high-molecular-weight poly-l-lysine (50 μg/mL; Sigma-Aldrich; A3890401), or high-molecular-weight hyaluronic acid (g/mL; Calbiochem), followed by rat tail collagen coating solution (Merck Life Science N.V.)(Hu *et al*, 2009, 2008) for 2h in incubator. Blocking was performed with 1% BSA in Dulbecco’s PBS (Life Technologies; 11500496)). After harvesting of cells, seeding of cells and incubation, plates were washed, fixed, and stained with Crystal Violet, and absorbance was measured at 590 nm as described. This protocol was replicated for three replicates per condition.

### Statistical Analysis

Data from in vitro experiments were presented as mean ± standard deviation (SD). Comparisons between two groups were conducted using a two-tailed Student’s t-test. For multiple group comparisons, we used one-way ANOVA followed by Bonferroni’s post hoc test to ascertain statistical significance. All analyses were performed using GraphPad Prism 9.5.1 software. Statistical significance was set at p < 0.05, with results denoted as *p < 0.05, **p < 0.01, ***p < 0.001. The Pearson correlation coefficient (r value) was calculated to evaluate linear relationships between variables. No predetermined method was employed for sample size determination, and normal distribution or variation of data was not assessed. In vivo study data were expressed as mean ± standard error (SEM), ensuring appropriate statistical representation. The analysis employed a chi-square test to explore the relationship between treatment groups and migration types in the study of zebrafish results. This statistical method evaluated the observed frequencies across the treatment conditions (Control and EFEMP1^KD^, or Control and EFEMP1^OEX^) and migration categories (0,1-4,5+).

### Public data analyses

The Breast Invasive Carcinoma dataset was obtained from TGCA using cBioPortal, a platform designed for the analysis of cancer genomics data (cBioPortal, n.d.; (Cerami *et al*, 2012). A search was conducted on EFEMP1 and STAT1 in cBioPortal to integrate clinical and gene expression data, providing valuable insights into the significance of these genes in breast cancer. The study also integrates information from the METABRIC dataset, which comprises gene expression data produced through microarray technology along with associated clinical data. This dataset was merged with the microarray expression data for additional analysis (cBioPortal, 2012).

To validate findings in publicly available human cancer cell line data, the Cancer Dependency Map (DepMap) platform was used. EFEMP1 and STAT1 expression data was compared in connection with the aneuploidy score.

### Data availability

RNA-sequencing data was uploaded to Array Express under accession number E-MTAB-14473.

## Acknowledgements

We are grateful to the members of the Foijer lab, van Vugt lab and other labs at ERIBA for fruitful discussion. Siqi Zheng and Ruifang Tian were supported by personal fellowships from the Chinese Scholar Council (CSC). This work was further supported by a Dutch Cancer Society and NWO Vici grant to Foijer (2015-RUG-7822 and 09150182210049). pLenti-pHluorin_M153R-CD63-mScarlet was a gift from Alissa Weaver (Addgene plasmid 172118; http://n2t.net/addgene:172118 ; RRID:Addgene_172118). We thank Nancy Halsema, Diana Spierings, Rianna Arjaans, Jennefer Beenen for help with RNA sequencing. We are grateful to Karina Köpke for help with imaging. We thank Ben Giepmans, Kim Kats and Anouk Wolters for assistance with TEM imaging. Mass spectrometry was performed at the Interfaculty Mass Spectrometry Center at RUG, with help of Marcel P. de Vries, Hjalmar Permentier and Dr. Karin Wolters. We thank Joop de Vries, Willem Woudstra, and Hélder A. Santos for providing access to nanoparticle quantification devices. Part of the work has been performed at the UMCG Imaging and Microscopy Center (UMIC), which is sponsored by the Netherlands Electron Microscopy Infrastructure (NEMI; NWO 184.034.014)

## Author contributions

SZ and RT performed most experiments, data processing, and analyses. KS supervised microscope imaging including TEM assays. JP performed zebrafish injections, and ED, and YL performed the confocal imaging of zebrafish embryos. RW, MS-PR, and PB assisted with experiments, LK performed RNA library preparation, RW analysed RNA sequencing data, MS analysed mass spectrometry data and performed public data analyses. EW and PvR assisted with nanoparticle characterization. MB helped optimizing EV assays under supervision of MD. PB engineered recombinant DNA construct design. SS co-supervised the project. SZ, and FF performed all project planning and experimental design. FF designed and supervised the study and provided funding. SZ and FF wrote the manuscript with input from all other co-authors.

## Supplementary Information

### Supplementary Tables

**Supplementary Table 1:** Complete mass spectrometry protein list.

**Supplementary Table 2:** The gRNA sequences, shRNA sequences, and QPCR sequences.

### Supplementary Movies

**Supplementary movie 1.** Time-lapse imaging movie demonstrating the uptake of pHluorin_M153R-labeled extracellular vesicles from BT549 donor cells to recipient BT549 cells, as confirmed in Supplementary Movie 1.

